# A single pathogen-secreted protein reprograms plants for drought resilience

**DOI:** 10.1101/2025.01.09.632073

**Authors:** Konrad Subieta, Leonie Weber, Jan-Niclas Lübbers, Karin Wiesotzki, Leon Pierdzig, Corinna Thurow, Johanna Schmitz, Ulrike Lipka, Kai Heimel, Kerstin Schmitt, Andrea Thürmer, Anja Poehlein, Rolf Daniel, Jan de Vries, Oliver Valerius, Gerhard H. Braus, Andrea Polle, Christiane Gatz, Thomas Teichmann, Volker Lipka

## Abstract

Climate change-enforced drought stress conditions and diseases caused by pathogens often co-occur and represent one of the greatest challenges in plant science^1–3^. Wilt pathogens that colonize water-conducting plant tissues can aggravate the problem and affect a wide range of agricultural crops^4,5^. However, whilst fungal infections with the vascular pathogen *Verticillium dahliae* are typically associated with wilt symptoms due to occlusion of xylem tissues, the related *V. longisporum* induces *de novo* formation of tracheary elements^6,7^. This promotes not only its virulence but also enables elevated water storage capacity of the infected host plant and resilience against drought stress conditions^6,7^. Here, we identified a secreted *Verticillium* protein, <u>TRA</u>NS<u>D</u>IFFERENTIATION <u>E</u>FFECTOR (TRADE), which triggers cell identity switches of bundle sheath cells into tracheary elements. We show that TRADE interacts with the intracellular plant protein VARICOSE (VCS), a conserved component of the mRNA turnover machinery and ortholog of the metazoan protein ENHANCER OF DECAPPING 4 (EDC4/HEDLS/Ge-1)^8^. The TRADE-VCS interaction induces SUCROSE NON-FERMENTING 1 (SNF1)-related protein kinase (SRK)-dependent phosphorylation and thus dysfunction of VCS. This affects the abundance of mRNAs encoding master regulators of xylem differentiation and demonstrates how a single pathogen effector protein triggers complex tissue-specific developmental reprogramming and thus promotes abiotic stress resilience.

---

In the course of evolution, plants have developed a variety of adaptive responses to abiotic and biotic stress conditions. This occurs at the level of reprogramming their transcriptome, proteome, metabolome and hormone balance, which ensures an adjustment of cellular processes, physiological homeostasis, growth, survival and reproduction^9,10^. Besides biotic stressors, such as microbial pathogens, insects, nematodes and parasitic plants, elevated temperatures and drought are the most widespread and climate change-enforced abiotic environmental stressors that affect the growth, development and productivity of plants^2^. Drought stress conditions and pathogen-induced diseases that often occur simultaneously under natural conditions pose a particular challenge, as they can act both, additively or antagonistically^1,3,4^. This applies especially to concurrent infections with fungal pathogens of vascular plants, including species of *Verticillium* and *Fusarium* that colonize the water-conducting tissues of their hosts and cause wilt disease^5^. Wilt symptoms are caused by vascular colonization, blockage of the water-bearing xylem vessels by fungal biomass and/or structural plant defence responses (tyloses) and fungal toxin production^11^. The most notorious crop pathogen within the genus *Verticillium* is the haploid species *V. dahliae*, which induces vascular wilt disease on over 200 plant species including tomato (*Solanum lycopersicum*), tobacco (*Nicotiana tabacum*), cotton (*Gossypium hirsutum*) and olive (*Olea europaea*)^11^. Another economically important *Verticillium* pathogen species is the allodiploid hybrid *V. longisporum*, which has a narrow natural host range and primarily infects dicot plants of the *Brassicaceae* family including rapeseed (*Brassica napus*), horseradish (*Armoracia rusticana*) and the model plant species *Arabidopsis thaliana*^12^. Despite proliferation in the vasculature, *V. longisporum* infections do not negatively affect the water status of their host plants and thus do not cause wilt phenotypes^13^. Instead, *V. longisporum* isolates induce chlorosis, stunting and vein clearing. Vein clearing is the consequence of cell identity switches of bundle sheath and xylem parenchyma cells into tracheary elements and the establishment of xylem hyperplasia^6,7^. This profound pathogen-induced developmental host plant reprogramming and *de novo* formation of water-conducting tissues is not only responsible for the lack of wilt symptoms, but even results in elevated host plant drought stress tolerance^6,7^. Cellular transdifferentiation triggered by *V. longisporum* involves the co-option of plant VASCULAR-RELATED NAC DOMAIN (VND) type NAC (for NAM, ATAF1/2, and CUC2) transcription factors^6,7^. However, the molecular mechanism by which the fungus employs these master regulators of xylem differentiation^14,15^ remains elusive.

To promote virulence and to create a suitable niche, fungal pathogens utilize host-specific effector molecules which interfere with the plant’s defence machinery, manipulate host microbiomes and/or reprogram host physiology, growth and development^16,17^. These effectors can be small molecules such as specialised metabolites, toxins, phytohormones and sRNAs, but also conventionally or unconventionally secreted proteins that target and act on extracellular or intracellular host proteins or processes^16–19^. Here, we report that a single lineage-specific *Verticillium* effector protein, TRANSDIFFERENTIATION EFFECTOR (TRADE), induces the described host cell identity switches by targeting a conserved scaffold protein of plant mRNA decapping complexes, VARICOSE (VCS), an ortholog of the metazoan protein EDC4/HEDLS/Ge-1^8,20,21^. TRADE-induced dysfunction of VCS alters transcript levels of central control elements of plant xylogenesis. Increased abundance of positive regulator transcripts, together with a decreased abundance of negative regulator transcripts mediates developmental reprogramming. Cell identity switches, *de novo* xylem formation and hyperplasia ultimately increase the plant’s water storage capacity and resistance to drought stress.

## *Verticillium*-triggered symptoms

To clarify whether the distinct interaction phenotypes are species-specific pathogen features, we conducted comparative infection experiments with a collection of 22 *V. longisporum* and 46 *V. dahliae* strains isolated from diverse dicot host plants and distinct geographical regions (Extended Data Table 1). Symptom development and disease index scoring identified eleven *V. longisporum* and three *V. dahliae* strains that did not induce any macroscopically detectable symptoms (Extended Data Table 1; Fig. 1a; Extended Data Fig. 1a). The remaining eleven *V. longisporum* isolates induced variable degrees of chlorosis and vein clearing disease symptoms on Arabidopsis Col-0. None of the tested *V. longisporum* strains induced wilt phenotypes, which was the prominent, but quantitatively variable disease symptom of the majority (i.e. 38) of *V. dahliae* strains. In agreement with Reusche et al.^7^, infections with wilt-inducing *V. dahliae* isolates did not trigger *de novo* xylem formation. Instead, enhanced lignification of xylem cell walls and limited adaxial expansion of vascular tissues were observed (Figure 1b, c, middle panel). Notably, five *V. dahliae* isolates (ST100, T9, V76, V138I and V781I) induced chloroses and cleared veins on Arabidopsis Col-0 (Extended Data Table 1; Fig. 1a; Extended Data Fig. 1a) concomitant with bundle sheath and xylem parenchyma cell transdifferentiation and establishment of xylem hyperplasia (Figure 1b, c, right panel; Extended Data Fig. 1b, c). Thus, the capacity to induce this developmental reprogramming pattern is not an exclusive feature of *V. longisporum*, but shared with a subset of *V. dahliae* isolates. Notably, all transdifferentiation-inducing *V. dahliae* strains were originally described as cotton or olive pathogens (Extended Data Table 1). Next, we compared the transdifferentiation-inducing *V. dahliae* and *V. longisporum* strains V76, V138I and VL43 with the wilt-inducing strain JR2 and another cotton-borne *V. dahliae* strain V192I, neither of which has the ability to induce host cell identity switches, in more detailed time-course infection analyses (Figures 1d, Extended Data Fig. 1d). This revealed that the transdifferentiation-inducing isolates were more virulent and that enhanced fungal proliferation correlated with reduced leaf size. Xylem hyperplasia induced by *V. longisporum* correlates with elevated transcript levels of the NAC domain transcription factor gene *VND7*^6,8^. Quantitative RT-PCR experiments confirmed a similarly increased transcript abundance for *VND7* and another NAC domain master switch of xylem formation, *VND2*^15^, in Arabidopsis infections with transdifferentiation-inducing, but not wilt-inducing *V. dahliae* strains (Fig. 1e; Extended Data Fig. 1e). Moreover, similar to *V. longisporum* VL43^6,7^, the cotton-borne transdifferentiation-inducing *V. dahliae* isolates V76 and V138I, but not the cotton strain V192I, induced enhanced drought stress tolerance in infected Arabidopsis plants (Fig. 1f, g; Extended Data Fig. 1f). This demonstrates that a subset of *V. dahliae* and *V. longisporum* isolates induce similar infection phenotypes in Arabidopsis. It is reasonable to assume that conserved and *in planta*-produced fungal effector molecules may be responsible.

**Fig. 1.**
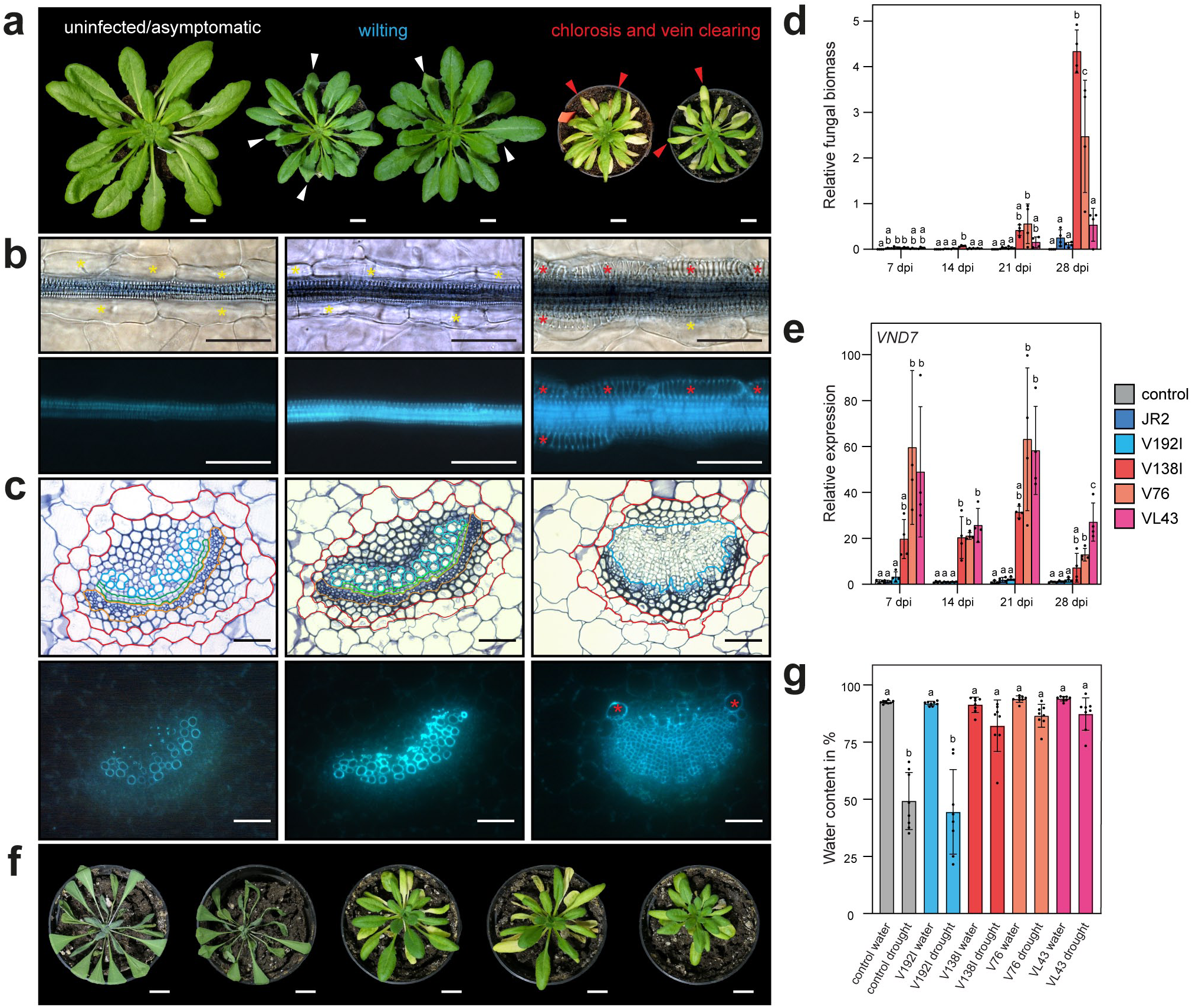
*V. longisporum* and a subset of *V. dahliae* share host plant infection phenotypes. **a**, Plant infection symptom classes “uninfected/asymptomatic”, “wilting” and “chlorosis & vein clearing”. Representative images from left to right: *V. longisporum* VL32, *V. dahliae* JR2, *V. dahliae* V192I, *V. longisporum* VL43 and *V. dahliae* V76 at 28 dpi. White arrows, wilting; red arrows, chlorosis & vein clearing. Scale bar, 1 cm. **b**, **c**, Representative images of leaf vasculature top view microscopy (**b**) and cross sections of leaf midrib veins (**c**). From left to right: *V. longisporum* VL32 (left), *V. dahliae* JR2 (middle) and V76 (right). Top, bright field microscopy; bottom, epifluorescence microscopy. Encircled tissues: red, bundle sheath cells; blue, xylem; green, cambium; ochre, phloem. Scale bar, 50 µm. **d**, **e**, *Verticillium* proliferation (**d**) and expression analysis of *VND7* (**e**) in leaves of control and infected plants. Bars represent means ± standard deviation (SD). Letters denote significant difference (one-way ANOVA with Tukey’s HSD p<0.05, n = 3-4 pools of 5 plants). **f**, Representative phenotypes of infected plants under restricted water supply at 24 dpi. From left to right; control, infected with *V. dahliae* V192I, V138I, V76 and *V. longisporum* VL43. Scale bar, 1 cm. **g**, Water content of plants under normal and restricted water supply in control and plants infected with *Verticillium* strains from (**f**) at 24 dpi. Bars represent means ± SD. Letters denote significant difference (one-way ANOVA with Tukey’s HSD p<0.05, n = 8 plants). Yellow asterisks, bundle sheath cells; red asterisks, bundle sheath cells transdifferentiated into xylem cells.

## Identification of *TRADE*

In transcriptome profiling experiments conducted with Arabidopsis root tissue infected by *V. longisporum* VL43, we identified *vl43-au17.g16224* as the most highly *in planta* expressed fungal gene in the VL43 genome^22^, whilst its expression was hardly detectable under *in vitro* growth conditions (Fig. 2a). The derived gene product is a protein of 237 amino acids with no known functional motifs, except for a predicted conventional N-terminal secretion signal peptide of 18 amino acids (Extended Data Fig. 2a). These features qualify the protein as a potentially secreted candidate molecule for the sought-after <u>TRA</u>NS<u>D</u>IFFERENTIATION <u>E</u>FFECTOR (TRADE). Interestingly, in the cotton *V. dahliae* isolate VD991^23^, a gene with very high sequence similarity is present on chromosome 4 in two identical copies. These two copies show opposite transcriptional directions in a duplicated, inverted genomic region of 39,485 nucleotides, which is separated by a unidirectional central spacer region of 341 nucleotides (Extended Data Fig. 2b-d). Southern blot experiments confirmed the presence of this tandem-inverted genomic repeat structure in all analysed transdifferentiation-inducing *V. dahliae* isolates and *V. longisporum* isolate VL43, but not in wilt-inducing strains (Figure 2b, c; Extended Data Fig. 2c, d), suggesting lineage-specific conservation. *TRADE* orthologs obtained from all transdifferentiation-inducing *V. dahliae* (*TRADE_Vd_*) and *V. longisporum* (*TRADE_Vl_*) isolates by PCR analyses showed a very high sequence identity of 99.3 % on the nucleotide and 98.7 % on the amino acid level between species and identical sequences within species (Fig. 2d; Extended Data Table 1, Extended Data Fig.1a). Transcript profiling in infected leaves of Arabidopsis and *N. benthamiana* corroborated the *in planta*-induced transcription of *TRADE_Vd_* and *TRADE_Vl_* (Fig. 2e, f). Consistently, chlorosis, but not wilt class pathotype *Verticillium* strains also induced developmental reprogramming and transdifferentiation patterns in *N. benthamiana* (Extended Data Fig. 2e). PCR amplification of the *TRADE* candidate gene failed in all isolates belonging to the wilt-inducing and asymptomatic class (Extended Data Table 1), supporting a causal relationship between presence-absence heterogeneity and pathotype. Notably, a single *TRADE-like* homolog is present in the genome of the wilt reference strain JR2^24^, with 90.3 % sequence identity on the nucleotide and only 81% identity on the predicted amino acid level to *TRADE_Vd_* (Fig. 2d; Extended Data Table 1; Extended Data Fig. 2a). We analysed all 68 *Verticillium* isolates used in this study and identified *TRADE-like* in all but the asymptomatic *V. longisporum* strains PD402 and PD730. The identified sequences can be assigned to five different *TRADE-like* alleles of *V. dahliae* and three different *TRADE-like* alleles of *V. longisporum* (Fig. 2d; Extended Data Fig. 2a). Notably, *TRADE-like_Vl1_*, *TRADE-like_Vl2_* and *TRADE-like_Vd5_*, that are present in transdifferentiation-inducing *V. dahliae* and *V. longisporum* isolates appear to be pseudogenes, suggesting a correlative lineage- and pathotype-specific *TRADE* gain and *TRADE-like* loss of gene function.

**Fig. 2.**
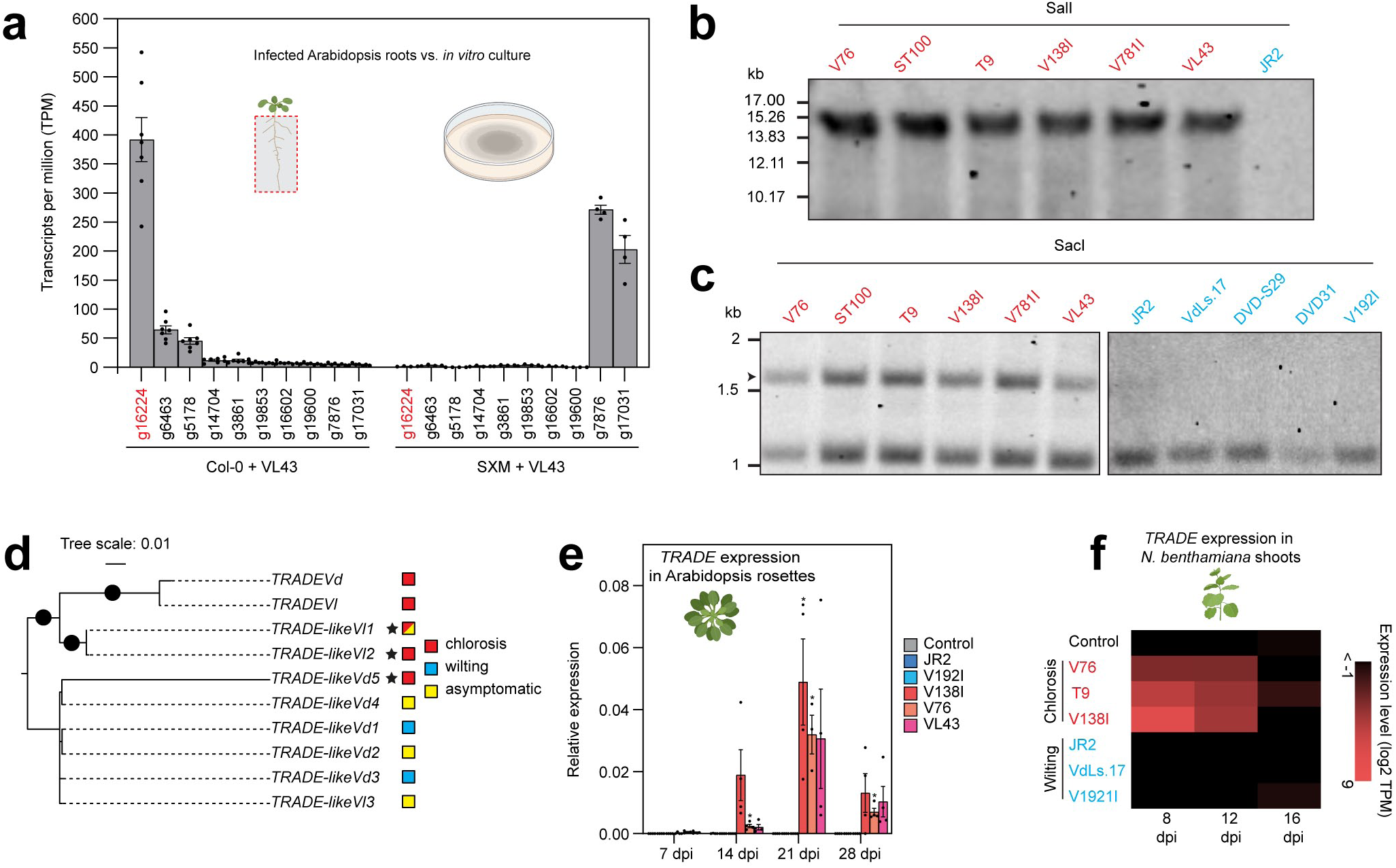
*V. longisporum* and *V. dahliae* chlorosis-inducing strains show high *in planta* expression of the conserved TRADE effector. **a**, Expression of fungal genes in transcripts per million (TPM) from *V. longisporum* VL43-infected Arabidopsis roots 10 days after infection (left) and 2 days growth on SXM agar plates (right). TPMs are relative to total transcripts found in each sample. Bars represent means ± standard error of the mean (SEM). **b**, **c**, Southern blots conducted on DNA of chlorosis- and wilting-inducing *Verticillium* strains digested with SalI (**b**) and SacI (**c**). Arrowhead in (**c**) indicates *TRADE* region specific bands. **d**, Maximum likelihood phylogeny of *TRADE* genomic nucleotide sequence homologs present in *V. dahliae* (*Vd*) and *V. longisporum* (*Vl*) isolates. Branches labelled with black dots indicate full ultrafast bootstrap (UFBoot) support. Stars indicate pseudogenes. **e**, Relative expression of *TRADE* in Arabidopsis rosettes of control and infected plants. Bars represent means ± SEM. Stars indicate significant differences to control within each time point. (Welch’s t-test, p<0.05, n = 3-4 pools of 5 plants). **f**, Heat map showing log_2_ TPM values for *TRADE* in control and infected *N. benthamiana* plants at 8, 12 and 16 dpi. Illustrations within **a**, **e** and **f** were created with Biorender.com.

## TRADE triggers host cell identity switches

To test if *TRADE* is required for virulence and encodes an effector protein with transdifferentiation capacity, both copies present in VL43 and V76 were deleted by homologous recombination (*ΔΔTRADE* lines; Fig. 3A; Extended Data Fig. 3a-f). As a complementary approach, one copy of the effector candidate gene driven by its native promoter was inserted into the genome of the wilt-inducing strain JR2 (Fig. 3A, Extended Data Fig. 3g, h). These genetic manipulations did not affect general fungal development and growth in axenic cultures (Extended Data Fig. 3i). Strikingly, Arabidopsis Col-0 plants did not show any macroscopic disease symptoms when inoculated with *ΔΔTRADE* lines (Fig. 3A). Epifluorescence microscopy confirmed that the bundle sheath cells remained intact and were indistinguishable from control plants (Fig. 3b, c; Extended Data Fig. 3j, k). In contrast, infection symptoms elicited by JR2 strains expressing TRADE were fully converted from wilt to chlorosis, phenocopying infections with V76 and VL43 at macroscopic and microscopic levels (Fig. 3a-c; Extended Data Fig. 3j, k). On the molecular level, deletion of *TRADE* in the V76 strain background abolished induction of *VND2* and *VND7* transcript levels, whereas expression of *TRADE* in the JR2 strain was sufficient to significantly increase transcript levels of both regulators of xylogenesis (Fig. 3d, e). Together, this validates the functional role of TRADE for the molecular reprogramming of host plant cell identity and development. To further address the molecular function of TRADE, we expressed Myc- and 6xHis-tagged TRADE_Vd_ and TRADE-like_Vd1_ without their own N-terminal signal peptide sequence in *Pichia pastoris* and purified the proteins by affinity chromatography. Upon infiltration into Arabidopsis rosette leaves, recombinant TRADE, but not TRADE-like, induced chlorosis and triggered transdifferentiation as well as xylem hyperplasia (Fig. 4a, b).

**Fig. 3.**
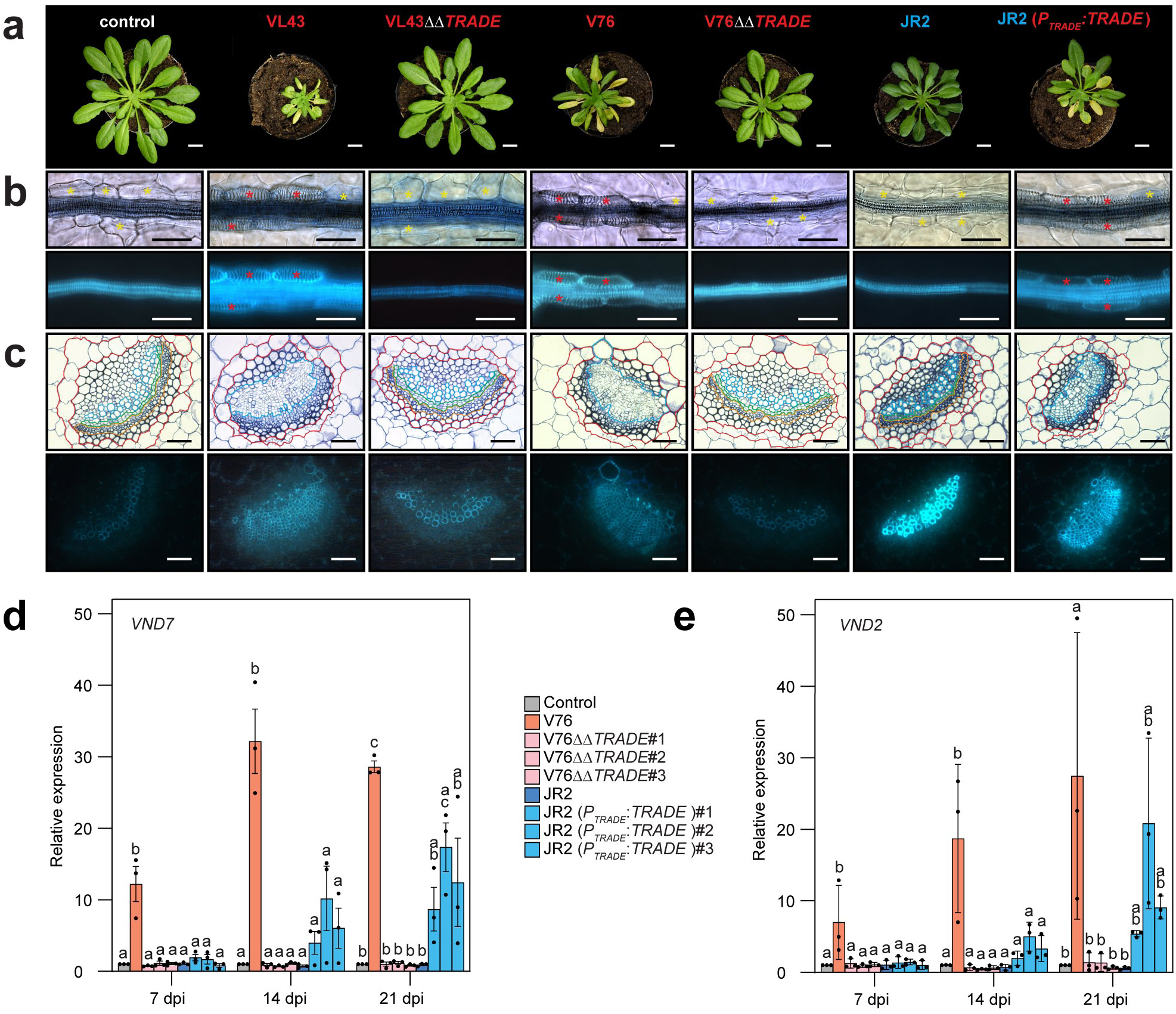
Transgenic *Verticillium* strains reveal that *TRADE* is necessary and sufficient for the establishment of chlorosis-type symptoms. **a,** Representative images of uninfected Arabidopsis control plants and plants infected with wildtype or transgenic *Verticillium* isolates at 21 dpi. *ΔΔTRADE*, *Verticillium TRADE* double knockout lines; JR2 (*P_TRADE_*:*TRADE*), JR2 *TRADE* expressing line. Scale bar, 1 cm. **b**, **c**, Representative images of leaf vasculature top view microscopy (**b**) and cross sections of leaf midrib veins (**c**) in infected plants. Top, bright field microscopy; bottom, epifluorescence microscopy. Encircled tissues: red, bundle sheath cells; blue, xylem; green, cambium; ochre, phloem. Scale bar, 50 µm. **d**, **e**, Expression analysis of *VND7* (**d**) and *VND2* (**e**) in leaves of control and infected plants. Bars represent means ± SD. Letters denote significant difference within the time points. (One-way ANOVA with Tukey’s HSD p<0.05, n = 3 pools of 4-5 plants from 3 independent experiments). Yellow asterisks, bundle sheath cells; red asterisks, bundle sheath cells transdifferentiated into xylem.

**Fig. 4.**
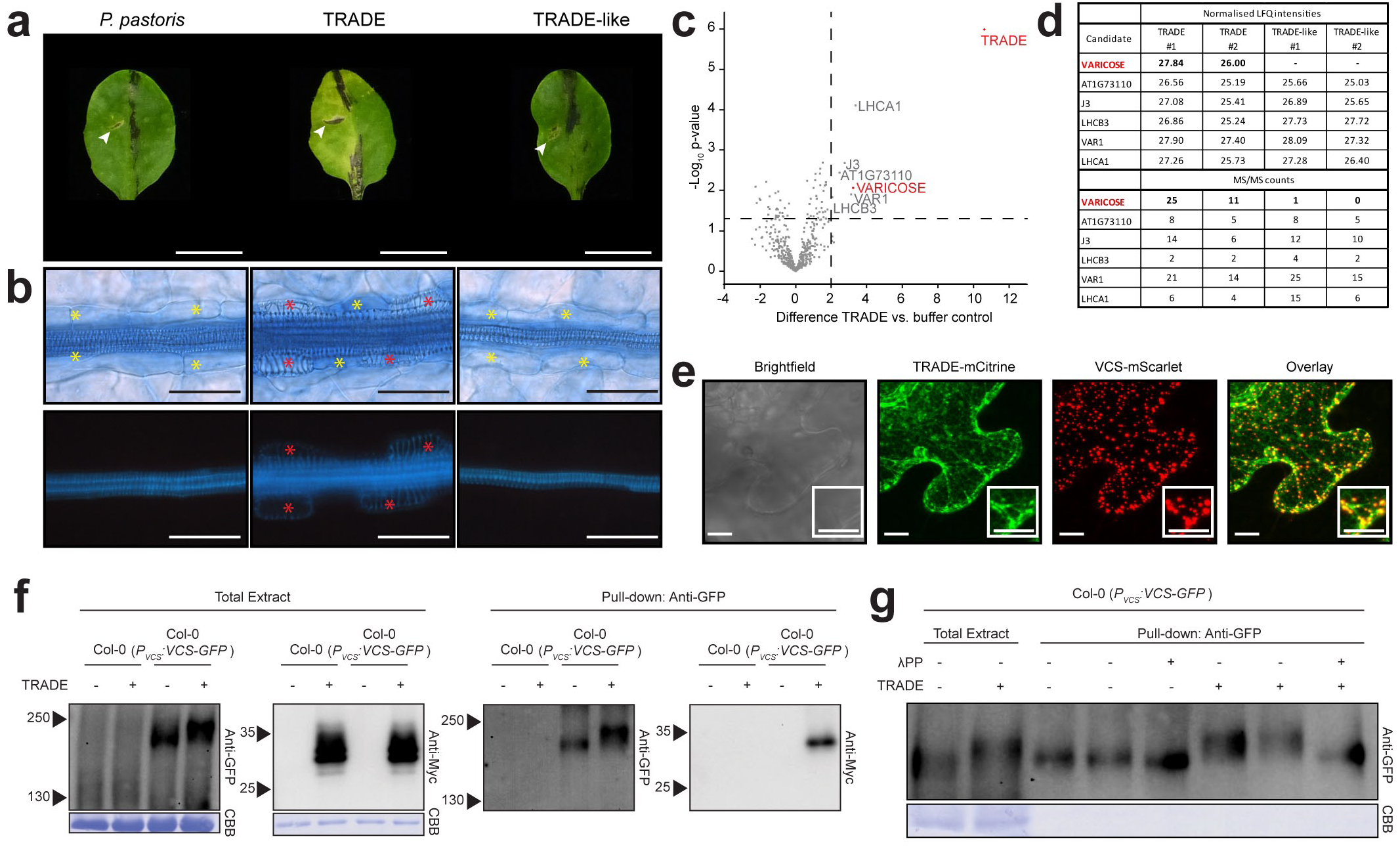
TRADE targets the mRNA decapping complex component VARICOSE (VCS) and induces its phosphorylation. **a**, Leaves of plants 7 days after infiltration with protein purified (left to right) from untransformed *P. pastoris* and from *TRADE* or *TRADE-like* expressing strains. White arrowheads mark infiltration sites. **b**, Leaf vasculature from leaves infiltrated in (**a**); upper panel, bright field microscopy; lower panel, epifluorescence microscopy. Yellow asterisks, bundle sheath cells; red asterisks, bundle sheath cells transdifferentiated into xylem. Scale bar, 50 µm. **c**, Volcano plot depicting TRADE interactors as determined via LC-MS analysis. X-axis; difference of the averaged log_2_ values of label free quantification (LFQ) intensities, dashed black line indicates cut-off at 2. Y-axis; significance values (-log_10_ p-value), dashed line indicates cut off at 1.3 (p<0.05). Data shown represent 4 independent experiments. **d**, Normalised LFQ intensities (top) and MS/MS counts (bottom) identified from two independent experiments in TRADE and TRADE-like infiltrated plants. Values shown for TRADE-treated samples originate from data also used in two of the four experiments in (**c**). **e**, Maximum z-projections of CLSM images from transient co-expression of *TRADE-mCitrine* and *VCS-mScarlet-I* in *N. benthamiana*. Leaf discs were infiltrated with 0.5 M NaCl to induce P-body formation. White boxes show an enlarged area of the respective images. Scale bar, 10 μm. **f**, Pull down experiment confirming the interaction of myc-tagged TRADE and VCS. **g**, Lambda phosphatase assay confirming TRADE-dependent VCS phosphorylation. CBB, coomassie brilliant blue.

To address TRADE localization and a potential intracellular function, we expressed a GFP-tagged TRADE variant lacking its N-terminal signal peptide in transgenic Arabidopsis plants under the control of the *UBIQUITIN10* (*UBQ10*) promoter. Phenotyping and gene expression analysis revealed variable transgene expression levels, which fully correlated with different degrees of stunted growth, macroscopically visible vein clearing and microscopically detectable transdifferentiation phenotypes (Extended Data Fig. 4a-c). Confocal laser scanning microscopy (CLSM) analyses with lines displaying intermediate expression levels showed ubiquitous expression and nucleo-cytosolic localization patterns in epidermal pavement cells, stomata and mesophyll cells (Extended Data Fig. 4d, e; Movie 1). Plasmolysis experiments revealed the absence of fluorescence signals in the plant apoplast (Extended Data Fig. 4f). Albeit the UBQ10 promoter confers tissue-independent expression, we observed particularly strong fluorescence associated with the leaf vasculature (Extended Data Fig. 4g). Intense TRADE-GFP signals were detectable in phloem companion cells and not yet transdifferentiated bundle sheath cells (Extended Data Fig. 4h), unravelling cell- and tissue-specific accumulation and localization patterns of the TRADE protein in the vasculature. Notably, fluorescence accumulated in (still) living leaf sheath bundle cells in granular structures (Extended Data Fig. 4i; Movie 2).

## TRADE targets VARICOSE

To identify *in planta* TRADE protein targets, we infiltrated purified TRADE and TRADE-like into Arabidopsis leaves. Subsequent pull downs and LC-MS analyses identified six proteins to be significantly enriched after TRADE treatment (Fig. 4c). Of these, only the VARICOSE (VCS) protein was specifically co-purified with TRADE, but not with TRADE-like (Fig. 4d). The other five potential interactors were present at similar levels in pull downs conducted with TRADE-like, suggesting that these proteins are not directly involved in the phenotypes induced by TRADE.

VCS is a key scaffold protein of the mRNA decapping complex, which is formed by physical interaction of VCS with the coactivator protein DECAPPING PROTEIN 1 (DCP1) and the catalytic subunit DCP2 to mediate 5’-3’ mRNA turnover in liquid-liquid phase separated cytosolic aggregates called processing (P) bodies^21^. The decapping complex is critical for early seedling development and normal vascular patterning, with mutants displaying seedling lethality and aberrant vascular tissues^20,21^. Indeed, comparative analyses of VCS and DCP2 loss-of-function mutants with *TRADE* expressing transgenic plants or seedlings treated with TRADE revealed strikingly similar developmental retardation phenotypes and ectopic xylem formation (Extended Data Fig. 4j, k). To test whether TRADE and VCS co-localize in P-bodies, we transiently co-expressed fluorescence-tagged fusion proteins in *Nicotiana benthamiana* leaves and induced P-body formation by NaCl treatment. We observed co-localization of TRADE-mCitrine and VCS-mScarlet in cytosolic aggregates (Fig. 4e). Together, these findings suggested that TRADE may target VCS in order to compromise its regulatory function in mRNA decay and homeostasis.

Reverse pull down experiments with TRADE-infiltrated Col-0 wildtype and transgenic plants expressing VCS-GFP under the control of its native promoter^25^ not only confirmed the physical VCS-TRADE protein interaction, but also revealed a TRADE-dependent mobility shift of the VCS protein (Fig. 4f). As VCS activity is regulated via phosphorylation^25^, we next performed lambda phosphatase treatments of pull downs isolated from control and TRADE-treated VCS-GFP expressing plants (Fig. 4g). The treatment reversed the TRADE-dependent VCS mobility shift, strongly suggesting that the TRADE-VCS interaction results in VCS phosphorylation and alteration of VCS functionality. Abscisic acid (ABA)-insensitive SNF1-Related Protein Kinases 2 (Class I SRK2s) SRK2A, B, G and H were previously shown to phosphorylate VCS in response to osmotic stress^25^. When we compared plants expressing VCS-GFP in Col-0 wildtype and *srk2abgh* quadruple mutant backgrounds in TRADE treatment experiments, *srk2abgh* mutant background plants were indistinguishable from the negative control, suggesting that they are insensitive to TRADE (Fig. 5a, b). Moreover, the phosphorylation-dependent mobility shift in response to TRADE treatment in Col-0 was fully absent in *srk2abgh* mutant plants (Fig. 5c). These results support the notion that the TRADE-induced phosphorylation of VCS requires ABA-insensitive SRK2 kinases and may be necessary for symptom development. Indeed, whilst Col-0 wildtype background plants infected with V76 developed typical, qualitative and quantitative chlorosis class symptoms, *srk2abgh* background plants showed no symptoms and markedly reduced fungal proliferation (Fig. 5d; Extended Data Fig. 5a-d). In infections with JR2, wilt phenotypes were observed in both tested plant backgrounds and fungal proliferation was largely unaffected. Hence, a general compatibility factor function of ABA-insensitive SRK2 kinases for *Verticillium* virulence can be excluded. Consistently, VCS phosphorylation was evident in the Col-0 background inoculated with V76, but not in the *srk2abgh* background or in extracts derived from JR2-inoculated plants (Fig. 5e). As V76 was unable to induce transdifferentiation in *srk2abgh* plants, we hypothesized that transcript accumulation of the VND master regulators of xylem differentiation upon infection would be abolished in this mutant line. Indeed, *VND7* and *VND2* expression was significantly lower in infected *srk2abgh* plants in comparison to plants in the Col-0 background (Extended Data Fig. 5e, f).

**Fig. 5.**
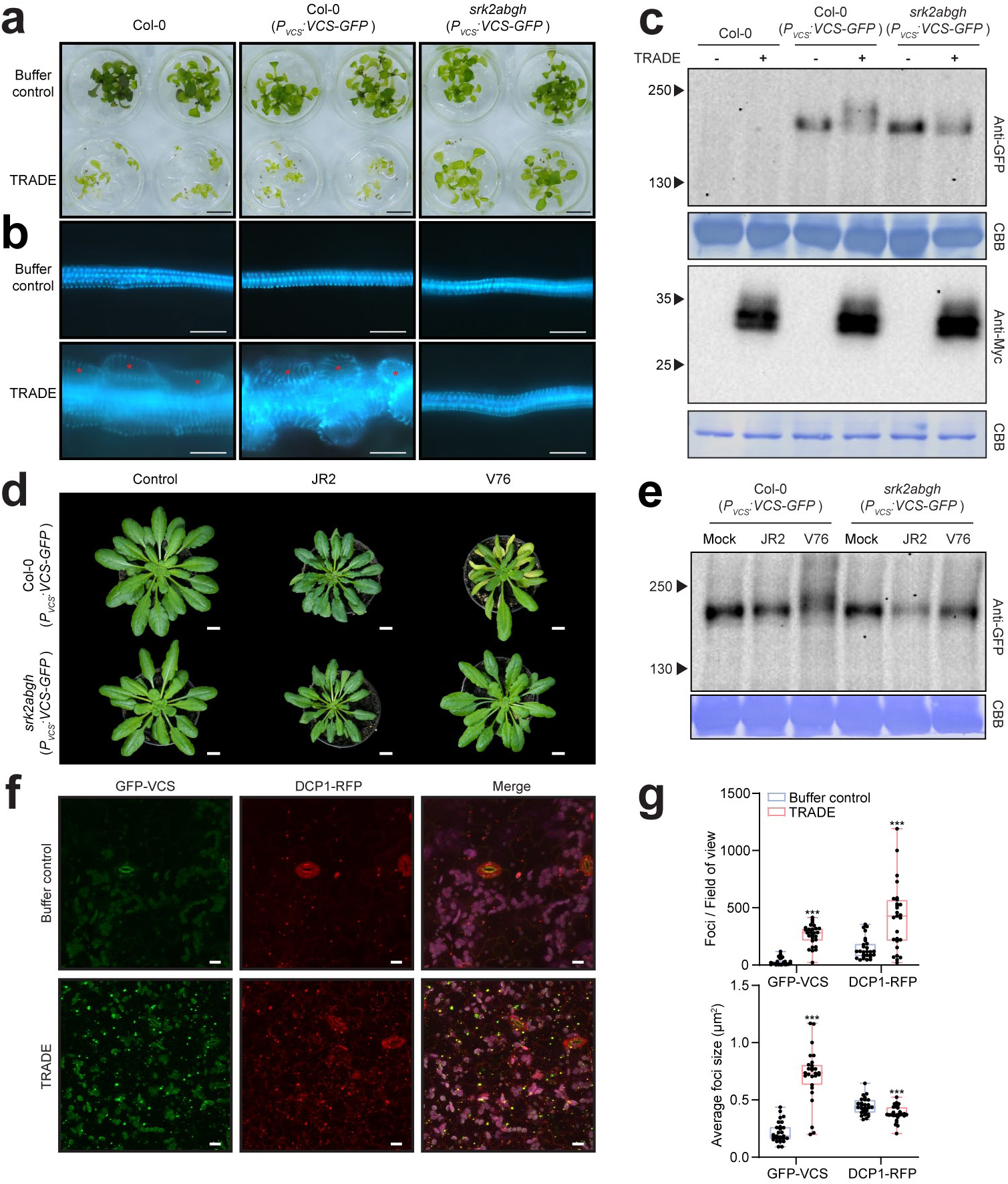
TRADE-associated phenotypes and VCS phosphorylation are abolished in the s*rk2abgh* mutant. **a**, TRADE treatment of wildtype and *VCS-GFP* expressing plants in the Col-0 and *srk2abgh* backgrounds in a well plate assay. Scale bar, 5 mm. **b**, Epifluorescence top view microscopy of leaf veins from plants in (**a**). Scale bar, 25 μm. **c**, Western blot of wildtype and *VCS-GFP* expressing plants in the Col-0 or *srk2abgh* backgrounds after TRADE treatment. **d**, Representative phenotypes of infected *VCS-GFP* expressing plants in the Col-0 and *srk2abgh* backgrounds at 28 dpi. Scale bar, 1 cm. **e**, Western blot of protein extracts from infected *VCS-GFP* expressing plants in the Col-0 and *srk2abgh* backgrounds. Plant extracts were prepared from a pool of five infected plants at 14 dpi. **f**, Maximum z-projections of CLSM images from plants expressing *GFP-VCS* and *DCP1-RFP* after buffer control and TRADE treatment. Chloroplast background fluorescence indicated as magenta in merged images. Scale bar, 10 μm. **g**, Quantification of foci per field of view (top) and average foci size (bottom) from plants treated in (**f**). Boxplot bounds represent the 25th and 75th percentile with the line showing the median. Error bars represent minimum and maximum values. Asterisks denote significant difference (n = 27, student’s t-test; ***, p<0.001).

## TRADE affects P-body behaviour

Stress inflicts changes in P-body size, number and protein composition^26,27^. These changes correlate with altered mRNA decay rates and were postulated to mediate the sequestration or release of translationally arrested mRNAs, allowing for rapid plant acclimation responses^28,29^. To address potential effects of TRADE on P-bodies, we performed CLSM analyses with leaves of double transgenic P-body marker lines expressing both GFP-VCS and DCP1-RFP^30^ (Fig. 5f). We detected significantly increased numbers of labeled foci per field of view for both GFP-VCS and DCP1-RFP upon TRADE treatment. However, while the average size of TRADE-induced VCS-GFP foci increased, this was not the case for DCP1-RFP tagged foci (Fig. 5g). Notably, TRADE-induced VCS aggregation was observed in both wildtype and *srk2abgh* mutant plants (Extended Data Fig. 5g, h). This suggests that the TRADE-VCS interaction results in the SRK2-independent sequestration of VCS, but not DCP1, into larger aggregates

## TRADE alters the plant transcriptome

Our data indicate that TRADE has two distinct effects on the VCS protein: a phosphorylation-independent sequestration into large aggregates and an SRK2-dependent phosphorylation, which may prevent the assembly of functional decapping complexes and thereby affect the levels of substrate mRNAs. To test this hypothesis, we carried out transcriptome time-course profiling analyses with Arabidopsis leaf material harvested at 0h, 6h, 12h, 24h, 48h, 72h and 96h after treatment with affinity-purified TRADE or the buffer as mock control. Principal component analysis (PCA) showed clear separation of control and treatment samples and clustering of biological replicates, except for separate clusters formed by the 6h and 12h samples, which may indicate infiltration-induced effects and/or circadian rhythm-dependent fluctuations (Extended Data Fig. 6a). Substantial numbers of differentially expressed genes (DEGs) were already detectable at 6h, with 1679 up- and 1302 down-regulated transcripts, which increased to >4400 up- and >5100 down-regulated transcripts in the plateau phase between 72h and 96h post-TRADE treatment (Extended Data Fig. 6b). Gene Ontology (GO) enrichment analyses revealed an over-representation of up-regulated terms associated with lignin, carbohydrate and cell wall biosynthesis as well as xylem/tracheary element development and differentiation (Extended Data Fig. 7). Terms enriched among down-regulated genes relate to pathogen defense and photosynthesis (Extended Data Fig. 8). Congruent with developmental reprogramming, cell identity switches and ectopic xylem formation, transcripts of genes with demonstrated roles in xylogenesis showed matching up- and down-regulation patterns (Fig. 6a). Up-regulated transcripts comprise key genes of all developmental phases of xylogenesis, ranging from procambial activity, initiation of xylem differentiation to secondary cell wall formation, and finally, programmed cell death^14,31–35^. The down-regulated transcripts include, in particular, genes that code for transcriptional suppressors of xylem differentiation^35^ and components of a signaling pathway that is crucial for vascular development and controlled by negative peptide regulators^36^.

**Fig. 6.**
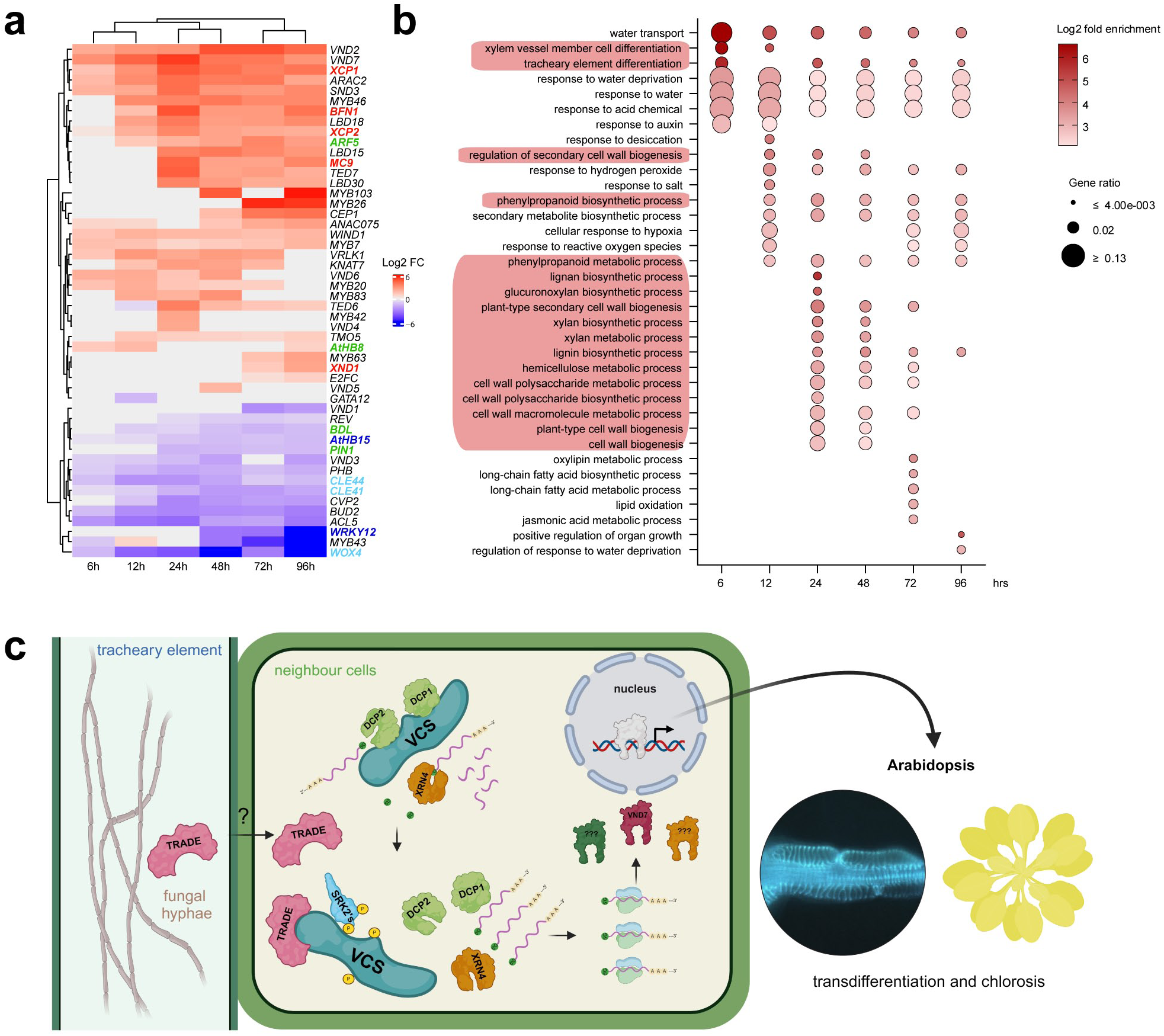
Accumulation of xylogenesis-related transcripts is similar in TRADE-treated and *vcs7* mutant plants. **a**, Heatmap showing DEGs related to xylogenesis after TRADE treatment (p<0.05). Red and blue bar colour codes indicate up- and down-regulation respectively, grey boxes indicate insignificant values. Gene acronyms coloured in green, (pro)cambial activity; black, xylem differentiation & secondary cell wall formation; red, programmed cell death; dark blue, transcriptional suppressors of xylem differentiation; light blue, vascular development controlled by negative peptide regulators. Fold change (FC). **b**, GO term enrichment analysis of upregulated DEGs overlapping between the TRADE transcriptome (log2 FC of ≥2 (padj<0.05)) and the *vcs7* mutant. Red background indicates GO terms involved in xylogenesis. **c**, Proposed model for TRADE activity. *Verticillium*-secreted TRADE is transported via an as of yet unknown mechanism into neighbour cells (i.e. bundle sheath or xylem parenchyma cells) of infected tracheary elements. Here, TRADE binds its target VARICOSE (VCS), inducing co-option of the ABA-independent SNF1-Related Protein Kinases (SRK2s) for VCS phosphorylation. Persistent phosphorylation leads to disruption of the decapping complex and aberrant mRNA turnover normally mediated by DCP1, DCP2 and XRN4. This ensues accumulation of a multitude of transcripts, including master regulators of xylem formation (such as *VND7*). This is responsible for the observed bundle sheath cell transdifferentiation and chlorosis in response to infection with *TRADE* harbouring fungi. Created with Biorender.com.

## TRADE and *vcs* transcriptomes overlap

Previously, Sorensen *et al.*^8^ used RNAseq and mathematical modelling to analyze mRNA turnover in Arabidopsis wildtype and mRNA decay mutants, including *vcs7*. This revealed complex feedback regulation of mRNA turnover and that VCS especially contributes to decay of short-lived RNAs, whose products are involved in procambium histogenesis, vasculature development and regulation of organ growth. To test our earlier conclusion that TRADE-dependent phosphorylation compromises VCS function and results in dysregulated mRNA turnover, we next compared DEGs identified in our transcriptome time course analysis to the VCS-dependent RNAs reported in Sorenson *et al*.^8^. GO term enrichment analyses of overlapping DEGs confirmed a significant over-representation of terms involved in xylogenesis (Fig. 6b). This further substantiates a mechanistic relationship between compromised VCS function, dysregulated mRNA turnover and ectopic xylem formation in both TRADE-treated wildtype and VCS loss-of-function mutant plants. Together with the GO term enrichment of photosynthesis-related processes reflected in the shared down-regulated transcripts (Extended Data Fig. 9) this provides a mechanistic framework to explain host plant chlorosis, stunted growth and other developmental phenotypes associated with TRADE effector secretion by transdifferentiation-inducing *Verticillium* isolates and *vcs* mutant plants.

## Discussion

This work suggests that the TRADE protein acts inside plant cells, which requires passage across host cell plasma membranes (Fig. 6c). Filamentous fungi likely deliver intracellular effectors into plant cells via clathrin-mediated endocytosis^37^, but the basidiomycete maize pathogen *Ustilago maydis* appears to produce a conserved set of proteins that were recently suggested to form a translocon structure for effector delivery^38^. Since yeast-expressed and affinity chromatography-purified TRADE protein alone is sufficient to induce cellular transdifferentiation, we can exclude the requirement of a fungal transfer machinery. Future genetic and pharmacological inhibitor experiments may either support the more likely scenario of endocytotic uptake or suggest a different, so far unknown transit mechanism.

The interaction with TRADE triggers VCS phosphorylation via the co-option of ABA-independent SRK2 kinases (Fig. 6c), which mediate rapid changes in mRNA levels by VCS phosphorylation in an osmotic stress response pathway conserved in land plants^25,39,40^. It has been suggested that the early and fast-acting ABA-independent osmotic stress response is post-translationally controlled by the ‘subclass I SRK2s-VARICOSE’ signaling module before ABA accumulation^39^. In this way, rapid ABA-independent signaling can be switched off (e.g. by dephosphorylation of VCS), while the ABA-dependent pathway takes control to prevent constitutively deregulated mRNA degradation required for normal growth and development. In contrast, TRADE-induced constitutive VCS phosphorylation can consequently lead to persistent dysregulation of 5’-mRNA decapping and turnover, affecting the abundance of transcripts with short half-lives, including mRNAs encoding regulators of xylogenesis (Fig. 6c;^8^). Even though we lack direct experimental evidence for a TRADE-induced change in the turnover of specific mRNAs encoding such key control elements, this conclusion is compelling, given the well documented and conserved function of the TRADE target protein VCS as a regulatory scaffold protein of decapping and mRNA turnover in plants and animals^8,20,21^ and the fact that not only the transcriptomes but also the phenotypes of *vcs7* knockout mutants and TRADE-treated plants are highly similar. Notably, the transcripts of *VND2* and *VND7* do not appear to be subject to direct VCS-dependent turnover^8^. This suggests control by higher-level, possibly cell-specific (transcriptional) regulators that are encoded by VCS-guarded mRNAs. We have reasonable confidence that such components can be obtained by isolating TRADE-insensitive plants from mutagenized Arabidopsis populations and subjecting them to molecular analysis.

TRADE-induced cell identity switches primarily affect vasculature-associated cells, such as bundle sheath and xylem parenchyma cells. In *Verticillium* infection experiments this may be explained by locally restricted TRADE production and secretion in colonized xylem cells, followed by diffusion and uptake into neighboring tissues. However, since infiltration of the purified protein and its stable transgenic expression also leads to restricted cellular protein accumulation and transdifferentiation patterns, TRADE-dependent effects are apparently related to cell type-specific reprogramming competence. This could be due to specific transcriptome signatures of vasculature-associated tissues, which may be characterized by a constitutive VCS-dependent turnover of transcripts encoding for regulators of xylogenesis. This is consistent with Sorensen *et al.*^8^, who proposed that spatial regulation of vascular development-related genes is impaired in *vcs* mutants and that VCS-mediated decay plays a role in defining tight tissue boundaries during development. Notably, Hirai *et al.*^41^ have recently shown that histone deacetylation controls xylem vessel cell differentiation and postulated that different patterns of histone deacetylase (HDAC) activity may be responsible for the different abilities of vasculature-associated cells and mesophyll cells to transdifferentiate either directly or with a prior dedifferentiation step. The application of TRADE in future HDAC inhibitor experiments and comparative single-cell RNA sequencing analyses may provide insights into the role of plant cell-specific DNA (de)acetylation, RNA homeostasis and turnover for plant stress response and developmental biology.

P-bodies are very dynamic super-assemblies of ribonucleoproteins (RNPs) undergoing phase transitions between soluble RNP complexes and condensed liquid or solid bodies in the cytosol. Their number, size, viscosity and demixing is tightly controlled by specific protein-RNA and protein-protein interactions, which also determine their function as sites of mRNA turnover or as storage compartments of repressed mRNAs that may reenter translation upon P-body dissolution^42^. Recent evidence suggests that large solid P-bodies are associated with mRNA storage, whereas small liquid ones are sites of mRNA degradation^43^. The idea that mRNA turnover is compartmentalized in small(er), potentially not even microscopically detectable RNP assemblies, is further supported by experiments showing that RNA degradation still occurs in the absence of visible P-bodies^44^. However, these general conclusions are still a matter of debate, recently fueled by a study showing that the functional role of P-bodies is modulated by cellular stress and that RNA decay occurs inside P-bodies and at faster decay rates^45^. Although TRADE treatment alters the size of P-bodies that contain VCS in Arabidopsis epidermis and mesophyll cells, TRADE-GFP only localizes to larger VCS-containing P-bodies in these cell types under osmotic stress conditions. Remarkably, TRADE-induced formation of larger VCS-specific P-bodies does not require SRK2-dependent phosphorylation. Synthesizing these results, we speculate that TRADE-binding induces conformational changes(s) in the VCS protein. This independently promotes VCS recruitment into larger P-body populations, as well as enhanced affinity of the VCS protein towards ABA-independent SRK2 kinases, ensuring durable VCS phosphorylation and dysfunction.

Given their function as an important site for controlled RNA decay and as a reservoir for rapid post-transcriptional adjustment of gene expression, it is not surprising that P-bodies have recently been discovered as hubs for plant immunity^46,47^ and pathogen targeting in both plants^30,48,49^ and mammals^50^. Our work now provides unprecedented insights and a mechanistic framework for how a single microbial effector molecule can induce intricate and fundamental developmental changes in their hosts via cell-type-specific modulation of mRNA homeostasis. This can be mutually beneficial under simultaneous abiotic stress conditions by providing a growth niche for the fungal invader while increasing the plant’s resistance to drought. The strong growth retardation associated with ubiquitous expression of *TRADE* in transgenic Arabidopsis plants so far precluded experiments to address the question whether plants producing TRADE exhibit enhanced drought stress resilience. Inducible, cell type-specific expression systems offer the opportunity for controlled temporal and spatial production of TRADE in xylem parenchyma, bundle sheath and other (control) cells. This will make it possible to test the biotechnological application potential to generate drought-resistant plants, one of the greatest challenges in plant biology posed by climate change^1–3^.

## Methods

### Plant material and growth conditions

The Arabidopsis ecotype Columbia-0 was used as a wildtype. Seeds grown on soil were stratified at 4 °C for two days in the dark and grown for 4-5 weeks under a 8 h light/16 h dark photoperiod at 22/18 °C, respectively. Light intensity and relative humidity were kept at 130 μE m^−2^ sec^−1^ and 65 % respectively. Wildtype *N. benthamiana* plants were grown for 4-5 weeks under a 16 h light/8 h dark photoperiod at 25/22 °C with the same light intensity and humidity as described above. Plants cultivated on ½ MS agar or in well plates were grown in a Percival growth chamber (Percival Scientific) under a 12 h light/12 h dark photoperiod at 22 °C with the same light intensity and humidity as described above. Published lines used in this study are: *P_VCS_:VCS-GFP* expressing plants in the Col-0 and *srk2abgh* background^25^, *GFP-VCS* and *DCP1-RFP* double marker line^30^. The T-DNA insertion mutant lines *vcs6* (SAIL_831_D08, N837130) and *dcp2-1* (SALK_000519, N500519) were obtained from the Nottingham Arabidopsis Stock Centre. The *TRADE-GFP* lines generated as a part of this study are in the Col-0 background.

### Generation of constructs

Primer sequences used can be found in Supplementary Table 1. In order to generate *TRADE* knockout constructs, LW125/LW139 and LW140/LW141 were used to amplify the 5’ and 3’ *TRADE* flanking regions from V76 genomic DNA respectively. The hygromycin cassette and vector backbone were amplified from *pPK2*^60^ with LW8/LW9 and LW10/LW11 respectively. The fragments were assembled using NEBuilder^®^ HiFi DNA Assembly (New England Biolabs) with the 5’ and 3’ regions flanking the hygromycin cassette, generating *pLW14*. For the nourseothricin cassette plasmid, LW125/LW189 and LW140/LW141 were used to amplify the 5’ and 3’ flanks, respectively. LW129/LW9 were used to amplify the nourseothricin cassette from *pSAB* (Department of Molecular Microbiology & Genetics, University of Göttingen, Germany; G.H. Braus). The vector backbone was amplified from *pPK2* using LW10/LW11. Fragments were assembled as described previously to generate *pLW19*. The plasmid used to express *TRADE* in JR2 was generated by amplifying the hygromycin and vector backbone from *pPK2* using LW8/LW10. The *TRADE* gene including 5’ and 3’ flanks was amplified from V76 genomic DNA using LW125/LW126. Fragments were assembled as previously mentioned to generate pLW11. To generate *TRADE-GFP* expressing lines, LW199/LW200 were used to amplify the *UBQ10* promoter and *GFP* containing vector backbone from *pHG113*. LW216/LW198 were used to amplify the *TRADE* gene without signal peptide and with the addition of a 4x alanine linker from V76 cDNA. The fragments were assembled as mentioned previously to generate pLW27. For expression of *TRADE* in *P. pastoris*, *TRADE* without signal peptide and with the addition of XbaI and EcoRI sites was amplified from V76 cDNA using LW281/LW280. The resulting fragment and the *pPICZα* vector backbone were digested using XbaI and EcoRI enzymes. Digestion products were ligated using the Rapid DNA Ligation Kit (Thermofisher Scientific) to yield pLW36. The *TRADE-like P. pastoris* expression plasmid was generated by amplifying the two exons of the *TRADE-like* gene without signal peptide from JR2 genomic DNA using LW281/LW284 and LW283/LW282. The exons were assembled using NEBuilder^®^ HiFi DNA Assembly, XbaI and EcoRI sites were then added to the resulting products by amplifying with LW281/LW282. The resulting products were then digested with XbaI and EcoRI alongside the *pPICZα* vector backbone and ligated as mentioned previously to yield pLW38. In order to generate the *TRADE-mCitrine* plasmid for transient expression, the *UBQ10* promoter and *TRADE* gene were amplified from pLW27 with oJL16/oJL17 and the vector backbone was a *pGreen-0229*^61^ plasmid containing *mCitrine* under the control of the *35S* promoter. Both fragments were digested with AscI and NotI and ligated as previously mentioned replacing the *35S* promoter for the *UBQ10* promoter and *TRADE* gene generating *pJL004*. To create the VCS plasmid for transient expression, the *VCS* gene was amplified from *pAMB122*^62^ with oJL018/oJL019 and the vector backbone was a *pGreen-0229*^61^ plasmid containing *mScarlet-I* under the control of the *35S* promoter. Both fragments were digested with SdaI and NotI and ligated as mentioned previously, inserting *VCS* between the *35S* promoter and *mScarlet-I* yielding *pJL005*.

### Generation of transgenic lines

Arabidopsis was transformed using the floral-dip method^63^. In order to transform *Verticillium*, a modified method for transformation of ascomycete spores was used^64^. A pre-culture was prepared by inoculating 2 mL of Luria-Bertani (LB) medium supplemented with antibiotics with one freshly grown *Agrobacterium tumefaciens* AGL1 colony containing the desired plasmid and incubating at 28 °C and 180 rpm for 20 hours. 1 mL of pre-culture was used to inoculate 20 mL induction medium^65^ in a 50 mL tube which was then incubated for 5 hours at 28 °C and 180 rpm. Afterwards, 500 μL induction medium containing *Agrobacteria* were mixed with 500 μL 1*10^6^ spores/mL *Verticillium* spp. spore suspension. From this suspension, 200 μL were pipetted onto solid induction medium plates (Ø 10 cm), covered with filter discs (Ø 85 mm, grade 3 hw, 65 g/m²; Sartorius, Göttingen, Germany). Plates were incubated at 25 °C in the dark for 3 days after which filter discs were transferred to Potato Dextrose Agar (PDA, Sigma-Aldrich) plates supplemented with cefotaxime as well as hygromycin or nourseothricin to select for transformed *Verticillium* spp. spores. Colonies were plated out twice on solid PDB containing appropriate antibiotics in order to avoid mixed cultures.

### Fungal culture

200 mL of simulated xylem medium (SXM)^66^ for spore production or potato dextrose broth (PDB, Sigma-Aldrich) for biomass production supplemented with 250 mg/L cefotaxime and 100 mg/L ampicillin were inoculated from a spore stock in 500 mL baffled Erlenmeyer flasks. Spore production was allowed to take place for 1 week in short day conditions (8 h light/16 h dark) on a rotary shaker at 90 rpm. Conidia were harvested by filtration through a folded filter (Macherey-Nagel MN 615 ¼; ⌀ 240 mm) and washed two times with sterile water. Spore concentration was determined with a Thoma counting chamber.

### Southern blot analysis of *Verticillium* spp. genomic DNA

DNA extraction was carried out as previously described^67^. 30 – 50 µg *Verticillium spp*. DNA was digested with restriction enzymes overnight. Fragments were separated on agarose gels and transferred by capillary blotting to an Amersham Hybond-N RPN203N membrane (GE Healthcare). DNA probes were generated with PCR and labelled using the Alkphos labelling module (RPN3680; GE Healthcare). 5’ and 3’ *TRADE* probes for digestions with BbsI and Bme1390I were generated with LW125/LW139 and LW140/LW141 respectively. 3’ probe for digestion with SalI was generated with LW243/LW215. The spacer region probe for digestion with SacI was generated with LW217/LW218. Hybridization of the probe was done following the protocol provided by the manufacturer (Alkphos labelling module, RPN3680; GE Healthcare). Star Detection Reagent (GE Healthcare) was used for probe detection and bioluminescence signals of the labelled probe were visualized in a ChemiDoc Touch (BioRad).

### *Verticillium* growth assay on solid medium

Growth dynamics of wildtype and transgenic *Verticillium* lines were tested by pipetting 5 µl of a 1*10^6^ spores/mL solution onto the centre of Czapek Dox agar plates supplemented with 250 mg/L cefotaxime and 100 mg/L ampicillin. Growth was monitored at 14 days after spore inoculation.

### *Verticillium* infection experiments

Infection experiments were carried out using an amended method^68^. Plants were grown in a 1:1 sand/soil mixture that was layered on granulated clay. The growth substrate was watered with 0.1 % plant fertilizer (Wuxal, Manna). Roots of three and a half week old Arabidopsis and three week old *N. benthamiana* plants were firstly wounded by gentle twirling, incubated in water for 30 minutes and subsequently incubated for 45 minutes in a conidial suspension of the respective *Verticillium* strain (1*10^6^ spores/mL), or in sterile water as a control. Plants were transplanted into single pots and kept under a transparent cover for two days to ensure high humidity. To monitor infection, Arabidopsis plants were photographed at 7 day intervals up to 21 or 28 days post inoculation (dpi). At each time point, whole rosettes of infected plants were harvested for DNA and RNA extraction. Leaf area was determined with the LeafScan software (Datinf GmbH).

### Determination of disease index

Infected plants were assigned to four infection classes: no symptoms (0), plants not stunted, but leaves chlorotic or plants slightly stunted without chlorotic leaves (1), plants moderately stunted and leaves chlorotic (2), plants severely stunted and many leaves chlorotic (3), plants dead (4). The disease index (DI) was calculated as follows: DI = ∑_i_ x j/n, where i is infection class, j is the number of plants in each class, and n is the total number of plants analysed^69^.

### Expression analyses and quantification of fungal biomass

RNA was extracted using Qiazol (Qiagen). The RNA pellet was re-suspended in ultra-pure ddH_2_O and remaining DNA was digested with DNase I, RNase free (Thermo Fisher Scientific). cDNA synthesis from 1 μg of total RNA was performed with the RevertAid™ H Minus First Strand cDNA Synthesis-Kit (Thermo Fisher Scientific). For fungal biomass quantification, total DNA was extracted with CTAB^70^. Realtime-PCR analyses were carried out in the C100 Touch thermocycler with a CFX96 system (Bio-Rad) using primers listed in (Supplemental Table 1). DNA and cDNA were used as template in reactions using the SsoFast EvaGreen mastermix (BioRad) for relative fungal biomass quantification and expression analyses respectively. Calculations for *VND2* and *VND7* expression were done according to the 2–ΔΔCT method^71^, with control plants being used for normalisation. For *TRADE* expression and fungal biomass quantification, 2–ΔCT was used^71^. The Arabidopsis *UBQ5* gene was used as reference.

### Drought stress experiments

At 14 dpi, all plants were watered until soil saturation. Subsequently, control plants were watered as normal, while watering of plants for drought stress experiments was stopped. Non-infected and infected plants were grown in separate trays on the same lighting rack. To avoid positional effects, plant trays were regularly turned 180° and the position of the trays in the climate chamber was regularly altered. Soil moisture levels were measured with a moisture meter (Acclima 305H-Sensor). To analyse plant water content, fresh weight and dry weight of rosettes was compared gravimetrically. For analysis of dry weight, plants were dried at 65 °C in an oven and dry weight was analysed immediately after drying. Water content was calculated as a percentage of plant fresh weight.

### *V. dahliae* VD991 *TRADE* region gene prediction

The 39,485bp *TRADE* containing region in the VD991 genome^23^ was searched for predicted genes by annotation from the published VL43 genome^22^ and by using the AUGUSTUS gene prediction tool with *V. longisporum* set as the organism parameter (https://bioinf.uni-greifswald.de/augustus/submission.php^72^). Coding sequences were generated by the removal of predicted introns and subsequently translated. Protein sequences were submitted to BLASTp, hits with the highest sequence identity were depicted in the scheme.

### Phylogenetic analyses of *TRADE* and *TRADE-like* genes

Primers specific for *TRADE* and *TRADE-like* (Supplementary Table 1) were used to amplify the respective genes in all 68 strains of a *Verticillium* collection (Extended Data Table 1). Identical *TRADE-like* sequences were assigned to the classes *Vl1* to *Vl3* (*V. longisporum*) and *Vd1* to *Vd5* (*Verticillium dahliae*). *V. longisporum* (*Vl*) and *Verticillium dahliae* (*Vd*) *TRADE* sequences and *TRADE-like* sequences of the eight classes were aligned using MAFFT v7.490^73^ with the option L-INS-I. A maximum likelihood tree was computed using IQ-Tree v. 1.5.5^74^ with the best fit model for sequence evolution determined by ModelFinder^75^; according to Bayesian Information Criterion; the best model was TPM3u. 1000 ultrafast bootstrap pseudo-replicates were computed^76^.

### Protein extraction from Arabidopsis

For proteomic analyses, Arabidopsis leaf material was ground on liquid nitrogen, 8 – 10 mL of powder was mixed with 10 mL of extraction buffer (250 mM sucrose, 5 % glycerol, 1 mM Na_2_MoO_4_ x 2 H_2_O, 25 mM NaF, 10 mM EDTA, 1 mM DTT, 0.5 % Triton X-100, 50 mM Na_4_P_2_O_7_ × 10 H_2_O, 100 mM HEPES-KOH pH 7.5 supplemented with 1:100 protease inhibitor cocktail (20 mM AEBSF, 72 mM bestatin hydrochloride, 72 mM pepstatin A, 1 mM leupeptin hemisulfate, 140 mM E-64, 277 mM phenanthroline)). Extracts were transferred into a 50 mL tube and centrifuged at 4000 rpm for 5 minutes at 4 °C. Protein concentrations were determined with Bradford reagent (Biorad) and subsequently equilibrated. For Western blotting, approximately 120 mg of pre-ground leaf material was placed in pre-cooled 1.5 mL microcentrifuge tubes with the addition of sand. After the addition of 200 μL of extraction buffer, the sample was immediately ground for 30-45 seconds using the IKA® RW20 digital drill (IKA-Werke) with a glass pistil at 1000 rpm. The pistil was promptly washed with 1 mL of extraction buffer and the liquid was collected into the micro centrifuge tube. The samples were centrifuged at 14.8k RPM for 6 minutes at 4 °C. The supernatant was transferred to a new 1.5 mL micro centrifuge tube. The protein concentration of each sample was determined with Bradford reagent, equilibrated and samples were mixed with 4x SDS loading dye.

### Western blot analyses

Proteins separated by SDS-PAGE were transferred to a PVDF membrane (Roti-PVDF, pore size 0.45 μm; Carl Roth) using electroblotting (Trans-Blot Cell, BioRad). For detection of VCS protein, the Pierce™ Western Blot Signal Enhancer (Thermofisher Scientific) was applied. After blocking membranes in TBS-T containing 3 % milk powder (w/v) primary antibody solution was added (α-GFP 3H9 or Myc-Tag 9B11) and membranes were incubated overnight in the dark at 4 °C. Membranes were washed and exposed to the secondary antibody (goat-anti-rat AP conjugate for α-GFP 3H9 and goat-anti-mouse AP conjugate for Myc-Tag 9B11) for 2 hours at RT. After final washing steps, Immun Star-AP substrate (BioRad) was added and signals were visualized using a ChemiDoc (BioRad).

### Protein expression in *P. pastoris*

The *P. pastoris* X33 strain was used for transformation and protein expression was performed using the EasySelect *Pichia* Expression Kit according to the manufacturer’s protocol (ThermofisherScientific, Invitrogen EasySelect manual). His-tagged protein was purified from the culture supernatant using a HisTrap™ FF Crude affinity chromatography column (Cytiva) on an Äkta purifier (Cytiva). Elution fractions containing protein as determined by SDS-Page were pooled in Vivaspin 20 (10k MWCO PES) (Sartorius) concentrators. Buffer exchange and desalting was performed with 10 mM HEPES. Protein concentration was measured with a Nanodrop OneC spectrophotometer (Thermofisher Scientific) using the predicted size and extinction coefficient parameters for the respective protein (TRADE; kDa: 27, Ex. Coeff: 41940; TRADE-Like; kDa: 26.86, Ex. Coeff: 45950).

### Treatment of plants with recombinant TRADE and TRADE-like protein

In order to monitor transdifferentiation, ethanol-sterilised Arabidopsis seeds were placed in 24 well plates (Sarstedt) in semi-solid media (½ MS, 0.2 % agarose) supplemented with TRADE protein to a final concentration of 0.2 mg/mL and allowed to grow for two weeks. Alternatively, plants were grown on soil and the 5^th^ leaf of 5 week old plants was infiltrated with 0.05 mg/mL TRADE or TRADE-like on the abaxial side with a needle-less syringe. For pull-down and interaction experiments, leaves of Arabidopsis were vacuum infiltrated with 0.2 mg/mL TRADE or TRADE-like using a desiccator with a vacuum pump. Leaves were then placed on moist filter paper within petri dishes sealed with parafilm and incubated for 4 hours. For all experiments, TRADE or TRADE-like protein was diluted with HEPES 10 mM. HEPES 10 mM solution without the addition of protein was used as a control throughout.

### Affinity bead pull-downs and on-bead trypsin digest for proteomic analyses

Affinity beads (Myc-Trap® Agarose, ChromoTek) were equilibrated with extraction buffer, added to the protein extracts in 15 mL tubes and placed in the dark at 4 °C. After 4 hours of incubation, the supernatant was removed and the beads were washed 3 times with 10 mL extraction buffer followed by three times with 10 mL wash buffer (150 mM NaCl, 0.5 mM EDTA, 10 mM Tris-HCl pH 7.5). Beads were then re-suspended in 1 mL wash buffer and transferred to 1.5 mL tubes, centrifuged and washed with 1 mL wash buffer. Beads were dried by extracting the excess wash buffer with a 0.5 mL U-100 insulin syringe (BD). For on-bead trypsin digestion, an adapted protocol from Chromotek was used. Beads were re-suspended in elution buffer 1 (2 M urea, 5 μg/mL trypsin, 1 mM DTT, 50 mM Tris-HCL pH 7.5), incubated at 30 °C for 90 minutes and centrifuged. The supernatant was transferred to a separate vial. Affinity beads were re-suspended in elution buffer 2 (2 M urea, 5 mM iodoacetamide, 50 mM Tris-HCl pH 7.5) and centrifuged. This step was repeated once and the supernatants of both elution steps were combined. The solution was incubated at 32 °C overnight. The reaction was stopped by the addition of 4,8 μL Trifluoroacetic acid (TFA). The tryptic digest was purified with a C18 stage tip (CDS Empore™, Thermo Fisher Scientific). Eluted peptide samples were dried using a SpeedVac (Eppendorf) at 45 °C. The dry pellet was suspended in proteomics sample buffer (2 % acetonitrile, 0.1 % formic acid) and incubated at 23 °C for 1 hour. Samples were sonicated twice, centrifuged and submitted for LC-MS analysis. Sample processing and data generation was carried out by the Service Unit LCMS Protein Analytics at the Institute of Microbiology and Genetics, Georg August-University Göttingen as reported previously^77^. Protein database searches and subsequent data analyses were carried out using MaxQuant (V.2.2.0.0) and Perseus (V.2.0.7.0) respectively.

### Proteomics data analysis

Unless explicitly stated, default parameters as defined in the Maxquant and Perseus software were used. Following data upload in Maxquant, label-free quantification was enabled under group-specific parameters. Under global parameters, the Araport 11 protein database^78^ with the addition of TRADE and TRADE-like proteins was uploaded. Under label free quantification (LFQ) options, iBAQ was enabled. The resulting data were uploaded to Perseus with LFQ intensities selected as the main parameter. The matrix was filtered to remove proteins only identified by site, reverse hits and potential contaminants. Following this, LFQ intensities were transformed (log2) and individual samples were grouped based on treatment. For statistical analysis, four independent experiments consisting of HEPES 10 mM and TRADE treatment were used. The matrix was filtered to only retain proteins that appear at least four times in at least one group. Following this, missing values were replaced from a normal distribution (width: 0.3, down shift: 1.8, mode: total matrix). A two-sample t-test was conducted on the matrix with the threshold p-value (0.05) being set for truncation and the threshold for the t-test difference being set as 2. The y and x axis line labels reflect these thresholds respectively. For comparisons of LFQ intensities and MS/MS counts between TRADE and TRADE-like, two independent experiments were used. The data from these two experiments were also used as part of the aforementioned statistical analysis. The matrix was filtered to only retain proteins that appear at least two times in one group, LFQ intensities and MS/MS counts were then compared for the candidates identified in the scatter plot analysis.

Raw data are deposited at: PXD050314 (www.ebi.ac.uk/pride/)

### Affinity bead pull-downs for analysis of TRADE-VCS interaction

GFP-Trap® Agarose beads (ChromoTek) were washed and pre-blocked in extraction buffer with 2 % BSA (w/v) for 2 hours at 4 °C. Protein extracts supplemented with 2 % BSA (w/v) were mixed with pre-blocked beads and incubated at 4 °C for 4 hours. After bead washing, bound proteins were eluted using the acid elution protocol from Chromotek. For Western blot analyses, aliquots were boiled at 98 °C for 3.5 minutes and run on 12 % gels in order to visualise TRADE with anti-myc antibody. Additional aliquots were not heat treated and run on 8 % gels in order to visualise VCS with anti-GFP antibody.

### De-phosphorylation assays

VCS-GFP was pulled down from total protein extracts of HEPES 10 mM and TRADE-treated plants with GFP-Trap® Agarose beads. After incubation and washing, beads were re-suspended in lambda phosphatase master mix (New England Biolabs) and split into three aliquots. Protein was immediately eluted from one aliquot and the sample was stored at −20 °C. λ phosphatase enzyme was added to one of the two remaining aliquots and both aliquots were incubated at 30 °C for 30 minutes. Protein was eluted from the beads using the acid elution protocol from Chromotek. Samples were subjected to Western blot analyses using anti-GFP antibody.

### Tissue embedding, staining and brightfield microscopy

Leaf midrib samples from Arabidopsis plants were fixed in 37 % formaldehyde, 100 % acetic acid and 70 % ethanol (FAE, 5:5:90, v/v/v) and dehydrated in an ethanol series of 50, 60, 70, 85, 95 and 100 % ethanol for 90 minutes per dehydration step. Samples were kept in 100 % EtOH overnight. On the next day, the EtOH was exchanged with fresh 100 % EtOH and samples were kept for two hours in the solution before further processing. Samples were embedded in cold curing resin based on hydroxyethylmethacrylate using Technovit 7100/Technovit 3040 (Kulzer) according to the protocol of the manufacturer. 5 µm sections were obtained with a microtome (Hyrax M55, Zeiss, Oberkochen, Germany) using a carbide knife. Sections were transferred to water-covered microscope slides and dried on a heating plate at 70 °C. For histochemical analyses, cross sections were stained with 0.05 % w/v toluidine blue for 1 minute. Trypan blue staining was carried out by submerging leaves in trypan blue solution (12.5 % (v/v) lactic acid, 12.5 % (v/v) glycerol, 12.5 % (v/v) phenol, 50 % (v/v) ethanol, 0.125 g/L trypan blue) and transferred to a boiling water bath for 1-5 minutes depending on the size of the leaves. The trypan blue solution was discarded and replaced with chloralhydrate solution (2.5 g/L) which was exchanged daily until the solution remained clear. Microscopy was carried out using a Leica DM 5000B microscope fitted with a CTR HS controller and DFC 300FX camera. Where necessary to account for brightness differences, images were adjusted using auto tone or level adjust tools with changes being applied to the whole image.

### Confocal laser scanning microscopy

Confocal Laser Scanning Microscopy was carried out with a Leica TCS SP5 or SP8 FALCON system. Excitation wave lengths of 488 nm for GFP, 514 nm for mCitrine and 561 nm for mScarletI/RFP were generated using an argon or DPSS 561 laser. Emission was captured with Leica HyD detectors between 500 and 540 nm for GFP, 580 and 620 nm for mScarletI/RFP and 525 and 560 nm for mCitrine. Chlorophyll auto-fluorescence was detected between 740 and 770 nm. Focal planes of z-stacks were maximum projected using Leica LAS AF (v.2.7.2). To monitor P-body behaviour after TRADE treatment, the abaxial sides of three leaves from three different plants were infiltrated with a syringe with 0.2 mg/mL TRADE or HEPES 10 mM as buffer control. Localisation changes were monitored 24 hours post infiltration. 21 µm deep (1 µm per layer) and 155 µm^2^ large z stacks were taken from the top of the abaxial side of infiltrated leaves. Quantifications were conducted with ImageJ 1.54f, using the Labkit Plugin^79^ using up to 27 images per line and condition.

### RNAseq analyses

For RNAseq comparisons between infected Arabidopsis roots and *in vitro* grown VL43, cultivation and inoculation of soil-grown plants as well as RNA extraction and mRNA sequencing were performed as described previously^80,81^. Ten days after infection, thoroughly washed roots of 12 single infected or mock-treated plants were combined for one replicate, respectively. RNAseq data from seven biological replicates for each treatment were obtained from plant material, which had been genotyped as wildtype^81^. For analysis of transcripts in *in vitro* grown VL43, SXM medium with the addition of 2 % agar was inoculated with a 1*10^5^ spores/mL solution of VL43. Growth was allowed in the dark for 2 days at 23 °C. Mycelium was collected from the plate surface using a scalpel and subsequently frozen in liquid nitrogen. For RNAseq, four biological replicates were analysed, each replicate obtained from one SXM agar plate. RNAseq data analyses were done using the Galaxy platform^82^. Mapping of reads to the combined Arabidopsis (TAIR10 release-51; https://ftp.ensemblgenomes.ebi.ac.uk/pub/plants/release-51/fasta/arabidopsis_thaliana/) and *V. longisporum* VL43 (vl43-au17; https://bioinf.uni-greifswald.de/bioinf/katharina/verticillium_full_data^22^) genome reference sequences and counting of the aligned exon reads were carried out with RNA STAR (Galaxy version 2.5.2b-2^83^). The transcripts per million (TPM) of each gene were calculated based on the gene length and the read count mapped to it in relation to all mapped read counts within a sample. For the ranking of fungal genes according to their *in planta* expression, we excluded those transcripts, which showed a similar number of mapped reads in RNAseq data obtained from both mock-treated and infected plants.

For RNAseq analysis in infected *N. benthamiana* plants, plants were infected as mentioned previously. Samples of aerial parts of *N. benthamiana* were collected at 8, 12 and 16 dpi and frozen in liquid nitrogen. RNA was extracted with TRIzol and mRNA sequencing was performed as described for infected Arabidopsis roots^80^. Reads were aligned to the combined *N. benthamiana* (Nbe_v1_scf.fa, https://nbenthamiana.jp^84^) and *V. dahliae* VD991 (https://www.ncbi.nlm.nih.gov/nucleotide/PRJNA939821^23^) genome reference sequences using RNA STAR (Galaxy version 2.5.2b-2^83^). Assembly of potential transcripts and quantification of gene counts were performed with StringTie (Galaxy Version 1.3.3.2^85^). TPM of each potential gene was calculated in relation to all mapped read counts.

For RNAseq analysis using purified TRADE protein, the abaxial side of the 5th leaf of 5 week old Arabidopsis plants was infiltrated with 0.05 mg/mL TRADE or 10 mM HEPES buffer using a needle-less syringe. 6 leaves were infiltrated per treatment per time point. RNA was extracted using Qiazol (Qiagen) as previously described. RNAseq analysis was processed by Novogene (GB). Differential expression analysis between TRADE-treated and control samples was performed using the DESeq2 R package. The resulting p values were adjusted using the Benjamini and Hochberg’s approach for controlling the False Discovery Rate (FDR). Heat map of up and downregulated genes in response to TRADE treatment was generated using the pheatmap and complex heatmap packages in R. Comparison of DEGs between TRADE treatment and the *vcs7* mutant relative to the wildtype Col-0 genotype was carried out in Excel. DEGs with a log2 fold change of ≥1 or ≤-1 respectively were extracted from the *vcs7* transcriptome (log2 fold change for the vcs sov/sov comparison^8^). These were compared to DEGs from each time point taken in response to TRADE treatment (DEGs with a log2 fold change of ≥2 or ≤-2 respectively). Genes found to be overlapping between the two datasets were searched for GO terms using the PANTHER overrepresentation test^86,87^ with a cut off of at least 5 fold enrichment.

Raw data are deposited at:

GSE269025 (www.ncbi.nlm.nih.gov/geo/query/acc.cgi?acc=GSE269025),
GSE269234 (www.ncbi.nlm.nih.gov/geo/query/acc.cgi?acc=GSE269234),
GSE269388 (www.ncbi.nlm.nih.gov/geo/query/acc.cgi?acc=GSE269388),

### Replication and statistical analysis

Infection experiments and associated quantitative data analyses were carried out at least 3 times with the exception of drought stress experiments, disease index scoring and infections on *P_VCS_(VCS-GFP)* expressing plants in the Col-0 and *srk2abgh* background which were carried out twice. Western blotting experiments were carried out three times with the exception of Lambda phosphatase treatment and VCS-GFP visualisation in infected plants, which were done twice. Qualitative and quantitative analyses of P-body behaviour after TRADE treatment were carried out 3 times. All other experiments were carried out at least twice with the exception of the following analyses: Southern blotting of *TRADE* region and double deletion strains, TRADE-GFP plasmolysis microscopy, and microscopy of *VCS-GFP* expressing plants in the Col-0 and *srk2abgh* background. All experimental repeats yielded similar results. Information related to statistical analyses and biological replicates is embedded in the figure legends. ANOVA with Tukey’s HSD and t-tests were carried out in RStudio and Microsoft Excel, respectively.

## Acknowledgements

We are grateful to Kazuko Yamaguchi-Shinozaki (University of Tokyo) for sharing the Col-0 (*P_VCS_:VCS-GFP*), *srk2abgh* (P_VCS_:VCS-GFP) and *srk2abgh* lines. We thank Anders Hafrén (Swedish University of Agricultural Sciences) for the *P_UBQ10_:GFP-VCS P_UBQ10_:DCP1-RFP* double marker line and Alexis Maizel (University of Heidelberg) for the pAMB122 plasmid. We acknowledge the support by Andreas von Tiedemann (University of Göttingen), Bart Thomma (University of Wageningen), and Rafael Jiménez-Díaz (University of Córdoba), who provided *V. longisporum* and *V. dahliae* isolates used in this study. We thank Sascha Dietrich (University of Göttingen) and Iker Irisarri (University of Göttingen) for support in bioinformatic sequence data analyses. We are grateful to Melanie Klenke for excellent technical support, and our team members for discussions and helpful advice. This research was funded by the Deutsche Forschungsgemeinschaft (DFG, German Research Foundation): LI1317/7-1 to V.L.; GRK 2172 to VL; INST 186/1277-1 FUGG (Confocal Microscope Leica TCS SP8); INST 186/1230-1 FUGG (Q Exactive HF/U3000); INST 186/1465-1 (position Ke.Sc.). K.S. was supported within the frame of the joint international DFG-IRTG/NSERC-CREATE training program IRTG 2172 “PRoTECT: Plant Responses To Eliminate Critical Threats” (GRK 2172, project number 273134146) at the Göttingen Graduate Center of Neurosciences, Biophysics, and Molecular Biosciences (GGNB). J.d.V. acknowledges support by the DFG-funded SPP 2237 “MAdLand” (528076711) and an ERC-Starting Grant under the European Union’s Horizon 2020 research and innovation programme (852725; “TerreStriAL”).

## Author contributions

V.L. conceived and conceptualized the study; K.S., L.W. and J.L. generated materials used in this study; K.S., K.W. and L.W. performed RT-qPCR and analysed the data; K.W., L.W., J.S. and K.S. performed *Verticillium* infection experiments; J.L., L.P., U.L., L.W., T.T. and K.S. conducted microscopy analyses; K.S. performed drought stress experiments; K.S., K.W., J.S, C.T., L.W., T.T., A.T., A.P. and J.d.V. performed sequence data and bioinformatic analyses; L.W. performed southern blot analyses; K.S. performed co-immunoprecipitation, western blots, pull-down experiments and dephosphorylation analyses; Ke.Sc. and O.V. performed LC-MS analyses; K.H., R.D., J.d.V., G.H.B., A.P., C.G. and T.T. analysed data, provided critical feedback and helped to shape the research; K.S., T.T. and V.L. wrote the original draft of the paper; K.S. and V.L. reviewed and edited the paper with input from all authors.

## Competing interests

The authors declare no competing interests.

## Additional information

### Supplementary information

The online version contains supplementary material (Movies 1 & 2; Supplementary Table 1). Sample

## Extended data figure/table legends

**Extended Data Table 1.**
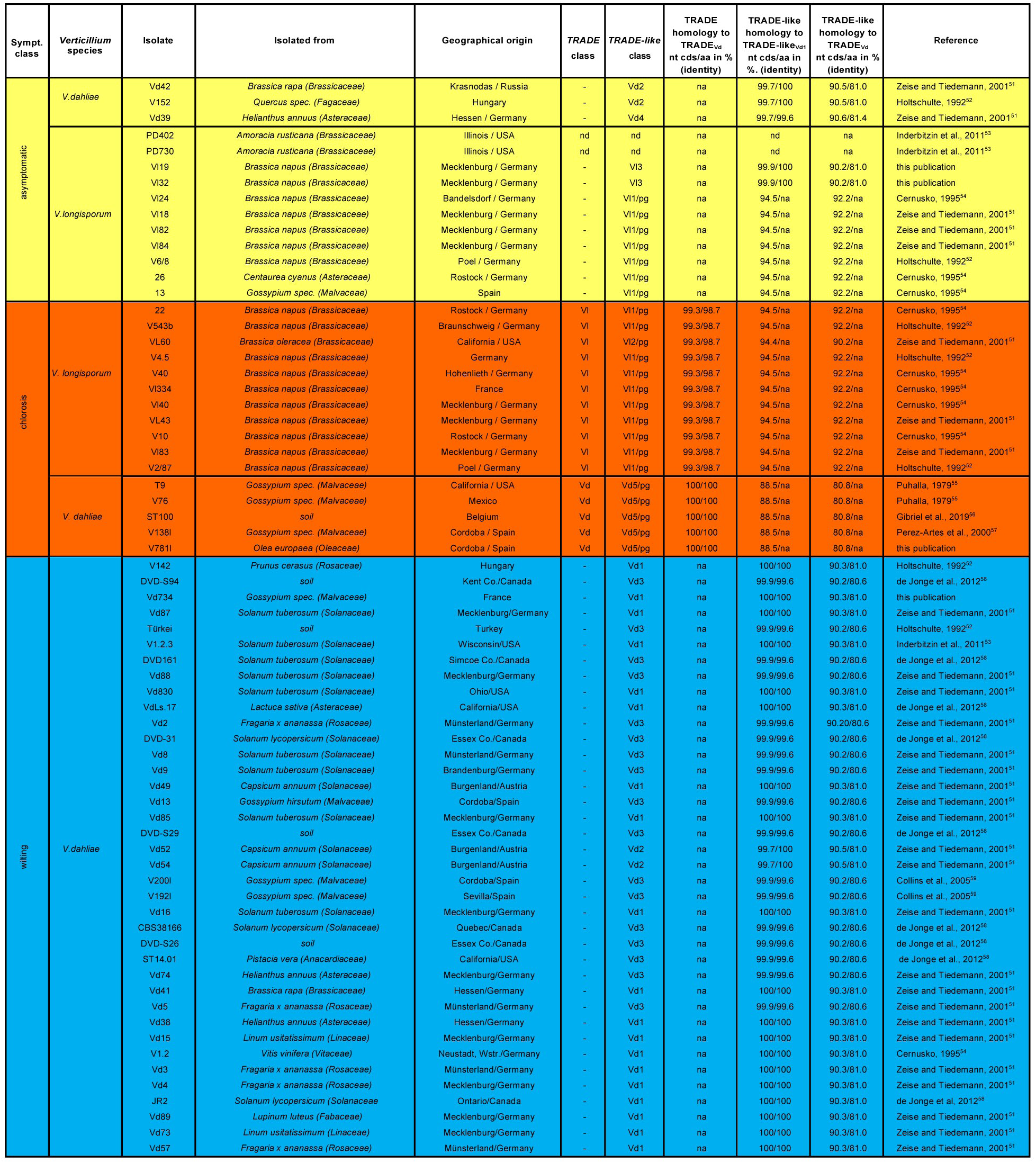
Presence-absence and sequence analysis of *TRADE* and *TRADE-like* in *Verticillium* strains used in this study. Colours indicate strain infection phenotypes as determined in Arabidopsis infections; yellow, asymptomatic; orange, chlorosis; blue, wilting. Sequence identity values on the nucleotide (coding sequence) and amino acid level calculated from sequenced *TRADE* and *TRADE-like* fragments. nd, not detected; na, not applicable; pg, pseudogene.

**Extended Data Fig. 1.**
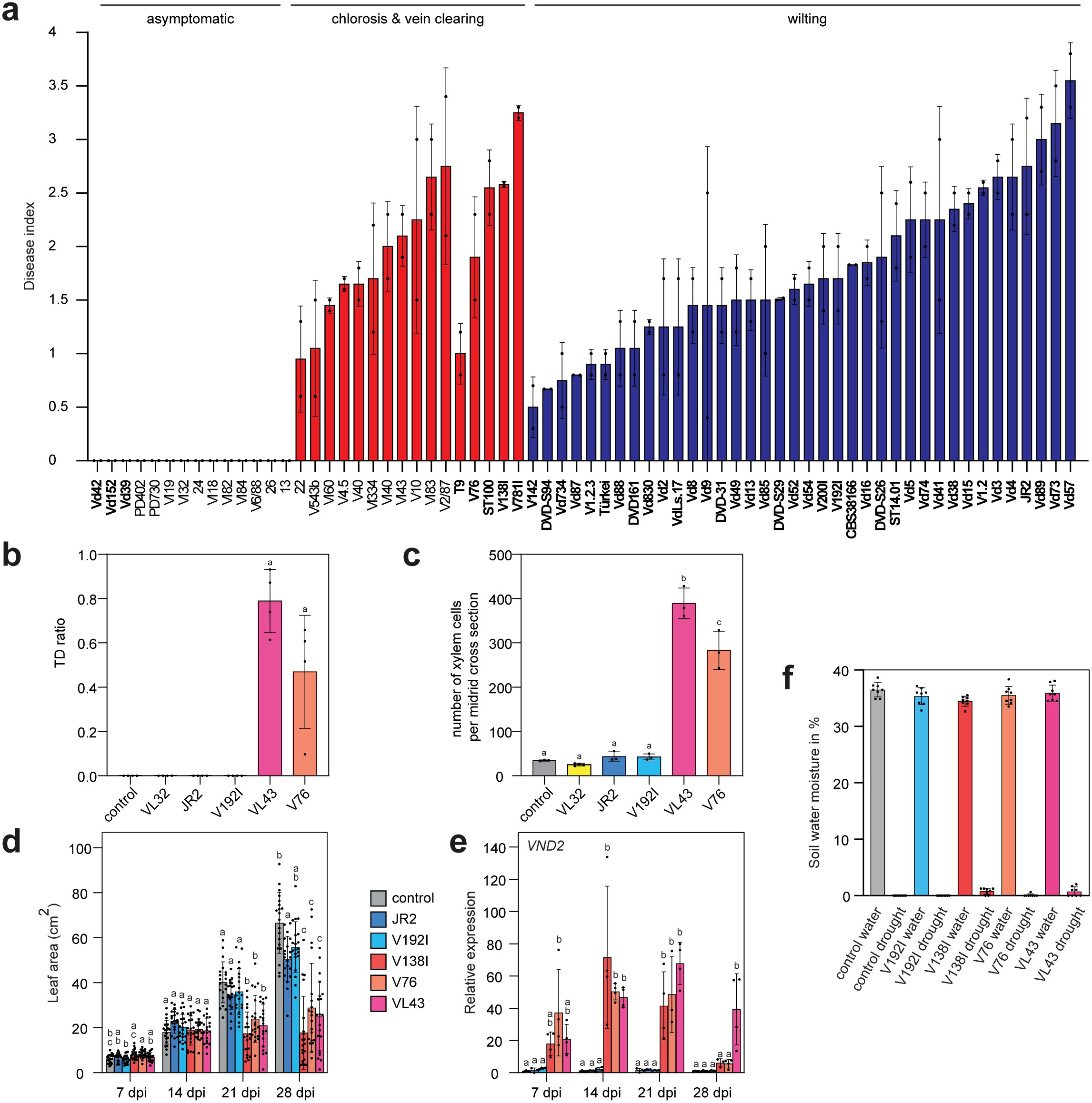
Quantitative analyses of *V. longisporum* and *V. dahliae* chlorosis and wilting strain-induced symptom development on Arabidopsis. **a**, Disease index of plants infected with asymptomatic, chlorosis and wilting strains. (n = 6-15 plants). Bars represent means ± SD calculated from 2 independent experiments. Red and blue colours represent chlorosis and wilting phenotypes respectively. *V. dahliae* isolates in bold characters. **b**, **c,** Quantitative analysis of transdifferentiation ratio in leaf vasculature top view microscopy (**b**) and number of xylem vessels per midrib cross section (**c**) in infected plants (referring to Fig. 1b,c). Bars represent means ± SEM. Letters denote significant difference. (One-way ANOVA with Tukey’s HSD p<0.05, n = 3-4). **d**, Leaf area of infected plants. Bars represent means ± SD. Letters denote significant difference within the time points. (One-way ANOVA with Tukey’s HSD p<0.05, n = 20 plants). **e**, Expression analysis of *VND2* in leaves of control and infected plants. Bars represent means ± SD. Letters denote significant difference within the time points. (One-way ANOVA with Tukey’s HSD p<0.05, n = 3-4 pools of 5 plants). **f**, Soil water moisture content in pots of control and infected plants grown under standard and restricted water supply. Bars represent means ± SD, (n = 8 pots).

**Extended Data Fig. 2.**
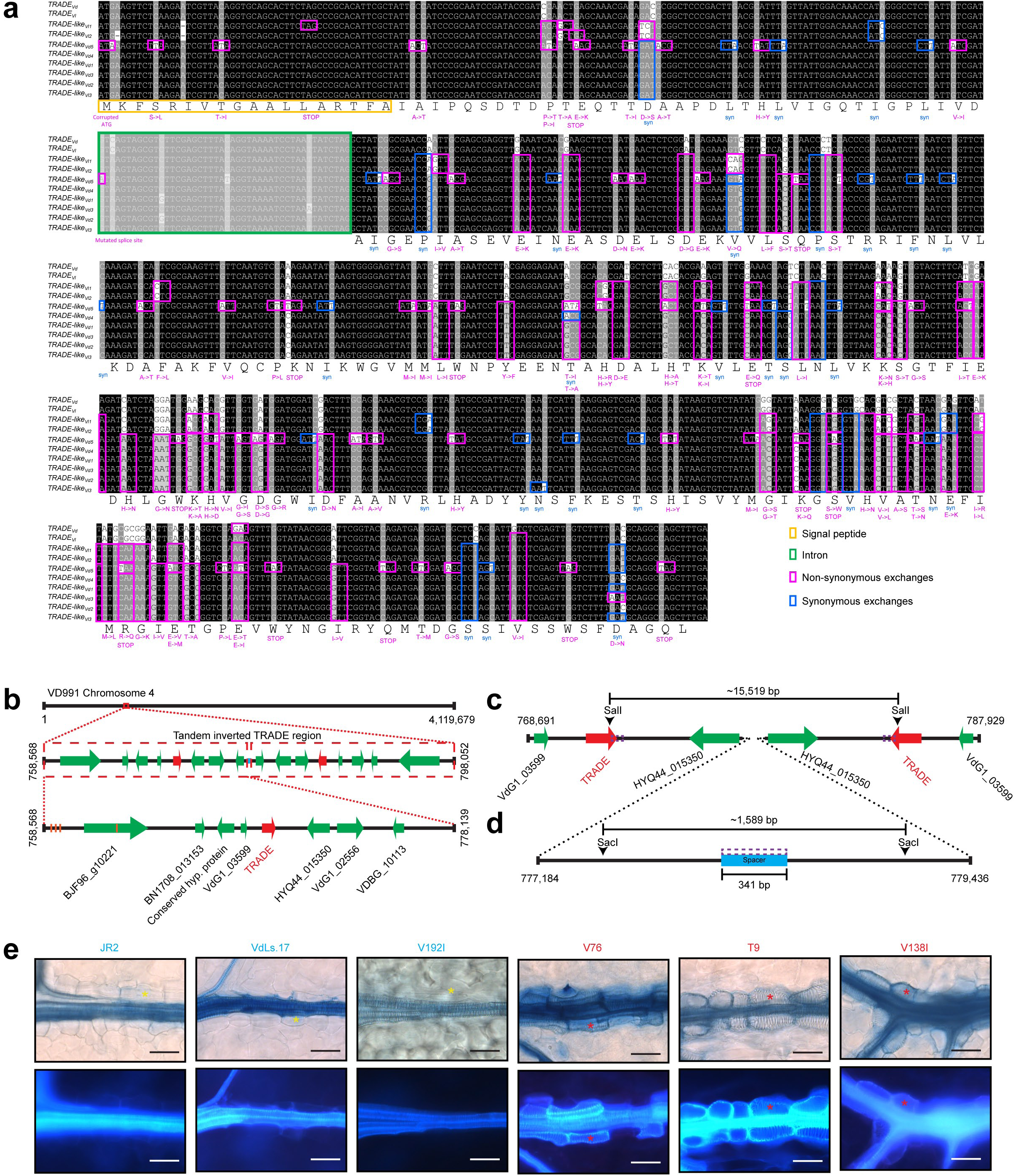
Nucleotide, amino acid sequences and genomic organization of TRADE variants, and transdifferentiation capacity of *V. dahliae* strains in *N. benthamiana.* **a**, Corresponding amino acid sequence of the reference *TRADE_Vd_* sequence shown below nucleotide alignment. Yellow box, signal peptide; green box, intron; blue boxes, synonymous exchanges; purple boxes, non-synonymous exchanges or mutated splice site. Exchanges are marked relative to the reference *TRADE_Vd_* sequence. Outcome of exchanges on the amino acid level is indicated in blue (synonymous) or purple (non-synonymous). Numbers at the end of alignment indicate length of nucleotide sequence for each *TRADE* and *TRADE-like* variant. **b**, Schematic representation of the tandem inverted *TRADE* region in *V. dahliae* VD991. Numbers left and right represent the base pair location of the depicted region. *TRADE* gene, red; other predicted genes, green; spacer, blue; SNPs, orange bars. Annotated gene labels based on highest homology in BLASTp searches. **c**, **d**, Schematic representation of restriction enzyme cleavage sites in the tandem inverted *TRADE* region (**c**) and spacer region (**d**) of *V. dahliae* VD991. Numbers left and right represent the base pair location of the depicted region. Cleavage sites are depicted with black arrows, bars represent predicted product sizes. Purple boxes represent *TRADE* 3’ (**c**) and spacer (**d**) probe binding regions. **e**, *N. benthamiana* phenotypes 16 dpi after infection with *V. dahliae* strains (wilt-inducing, blue; chlorosis-inducing, red). Bright field (top) and epifluorescence (bottom) microscopy images are shown. Scale bar, 50 µm. Yellow asterisks, bundle sheath cells; red asterisks, bundle sheath cells transdifferentiated into xylem cells.

**Extended Data Fig. 3.**
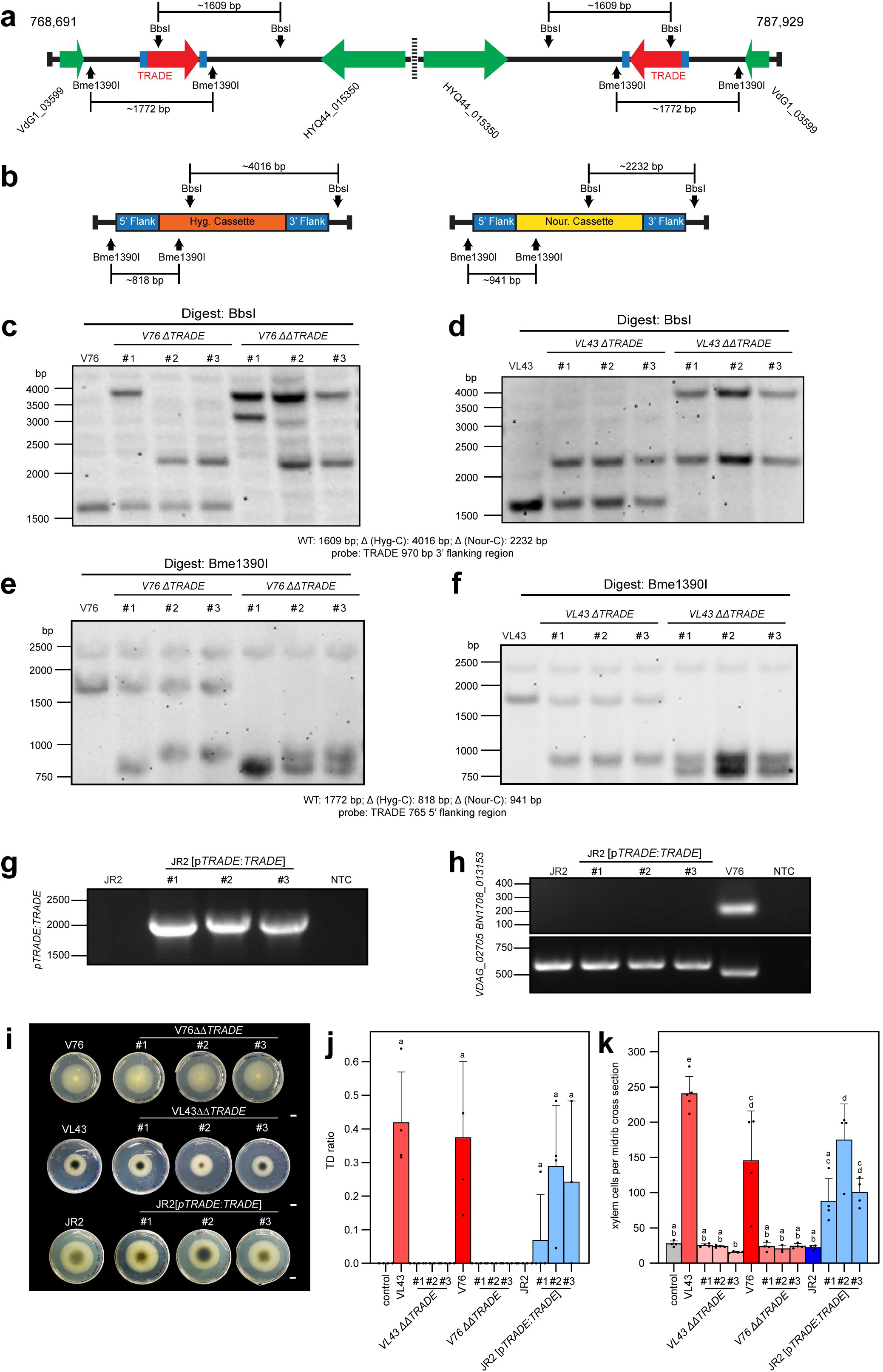
Double knockout and ectopic expression of *TRADE* does not perturb growth in axenic cultures. **a**, Schematic representation of restriction enzyme cleavage sites in the tandem inverted *TRADE* region of *V. dahliae* VD991. Numbers top left and right represent the base pair location of the depicted region. Cleavage sites are depicted with black arrows, bars represent predicted product sizes. Blue boxes adjacent to *TRADE* indicate 5’ and 3’ flanks used for homologous recombination with resistance cassettes. **b**, Scheme representing Hygromycin (left) and Nourseothricin (right) resistance cassettes used to generate *TRADE* knockout *Verticillium* strains. 5’ and 3’ flanking regions are shown as blue boxes. Restriction enzyme cleavage sites are depicted with black arrows, bars represent the predicted product sizes. **c**-**f**, Southern blots of wild type *V. dahliae* V76, *V. longisporum* VL43, single (Δ) and double (ΔΔ) *TRADE* knockout line DNA digested with BbsI (**c**, **d**) or Bme1390I (**e**, **f**). *TRADE* 3’ and 5’ flanking regions were used as probes for southern blot analysis in (**c**, **d**) and (**e**, **f**) respectively. **g**, **h**, PCR on genomic DNA from wildtype *V. dahliae* JR2 and three independent *TRADE* expressing JR2 lines confirming the presence of the *TRADE* expression construct (**g**) and absence of contaminating V76 DNA (**h**) in JR2 (*P_TRADE_:TRADE*) lines. The gene encoding BN1708_013153 was used to detect the presence of the *TRADE* genomic region. Gene *VDAG_02705* was used as a positive control for the presence of *Verticillium* DNA. **i**, Growth phenotypes of *V. dahliae* V76 and V76 *TRADE* double (ΔΔ) knockout lines (top), *V. longisporum* VL43 and VL43 *TRADE* double (ΔΔ) knockout lines (middle) and *V. dahliae* JR2, and JR2 (*P_TRADE_:TRADE*) lines (bottom) on Czapek Dox medium. Colony growth was monitored after 14 days of growth. Scale bar, 1 cm. **j**, **k,** Quantitative analysis of transdifferentiation ratio in leaf vasculature top view microscopy (**j**) and number of xylem vessels per midrib cross section (**k**) in infected plants (referring to Fig. 3b,c). Bars represent means + SEM. Letters denote significant difference. (One-way ANOVA with Tukey’s HSD p<0.05, n= 3-4).

**Extended Data Fig. 4.**
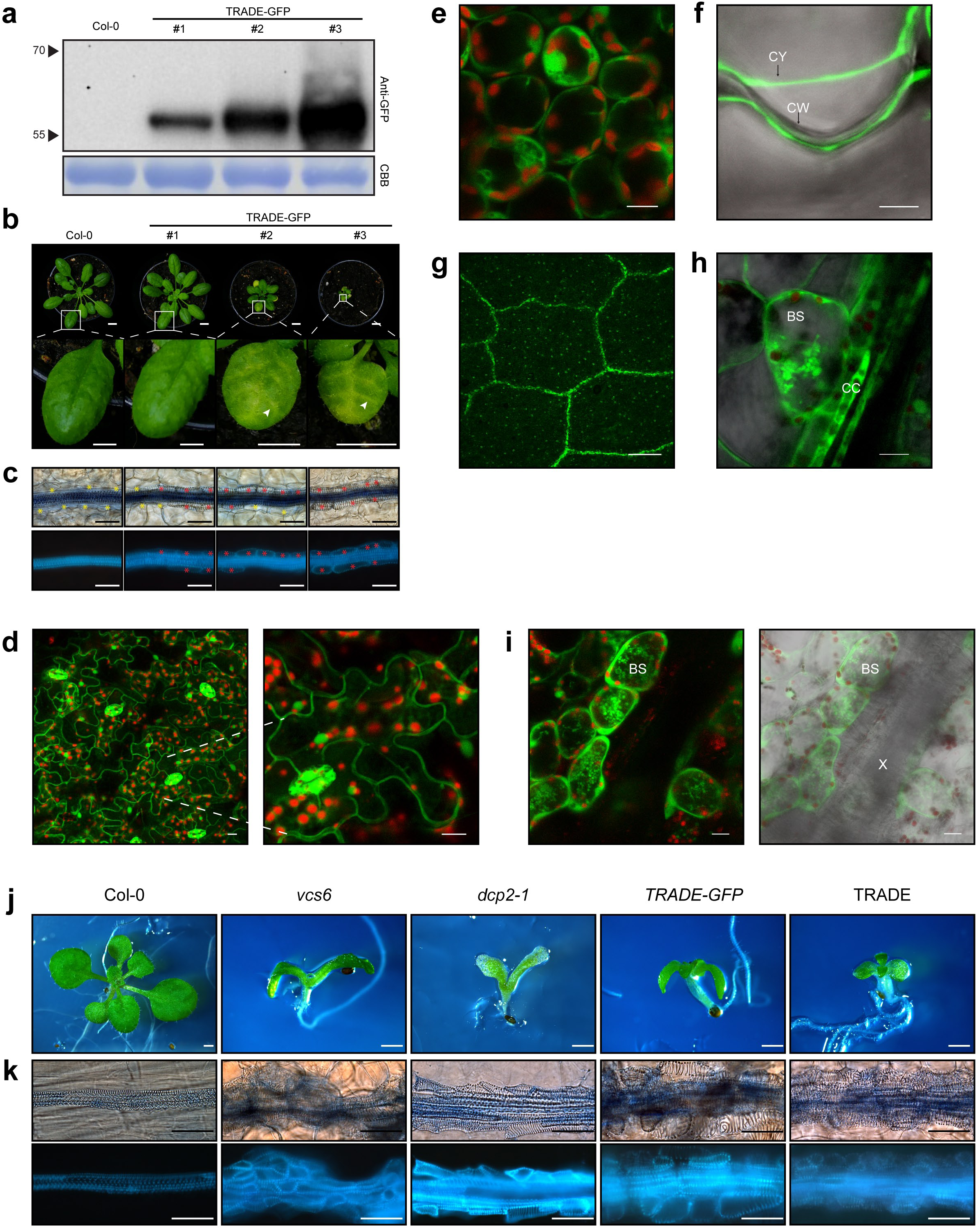
The TRADE protein alone triggers transdifferentiation and phenotypes similar to decapping complex mutants. **a**, Total protein extracts from wildtype and *TRADE-GFP* expressing plants. Detection with anti-GFP antibody. Bottom panel: coomassie brilliant blue (CBB). **b**, Images of wild type and *TRADE-GFP* expressing plants used in (**a**) (top row), scale bar, 1 cm. Enlarged view of leaf phenotypes (bottom row). White arrowheads indicate vein clearing. Scale bar, 5 mm. **c**, Microscopic images showing leaf vasculature of wildtype and TRADE-GFP expressing lines. Upper row, bright field microscopy; bottom row, epifluorescence microscopy. Scale bar, 50 µm. **d**, Representative maximum z-projections of CLSM images showing the subcellular localization of TRADE-GFP in leaf epidermis cells (left). Enlargement of area indicated by white dashed line (right). Scale bar, 10 µm. **e**, Representative CLSM images of mesophyll cells in *TRADE-GFP* expressing plants showing nucleo-cytosolic localisation. Overlay of chloroplast auto fluorescence with GFP channel is shown. Scale bar, 10 µm. **f**, Representative CLSM image confirming the absence of TRADE-GFP signals in the apoplast. Leaf disc treatment with 0.8 M mannitol for 30 minutes. CY, cytoplasm; CW, cell wall. Overlay of the bright field and GFP channels. Scale bar, 5 µm. **g**, Representative CLSM images showing TRADE-GFP signals within the leaf vasculature. Scale bar, 250 µm. **h**, Representative CLSM image showing TRADE-GFP signals within phloem companion cells (CC) as well as TRADE-GFP granular structures within bundle sheath cells (BS) of *TRADE-GFP* expressing plants. Overlay of the bright field and GFP channels. Scale bar, 10 µm. **i**, Representative maximum z-projections of CLSM images of xylem and associated bundle sheath cells in *TRADE-GFP* expressing plants (left), image overlay with bright field channel (right). BS, bundle sheath cell; X, xylem. Scale bar, 10 µm. **j**, **k**, Representative macroscopic and microscopic phenotypes of (left to right) wildtype, *vcs6*, *dcp2-1*, and *TRADE-GFP* expressing plants grown on ½ MS plant medium. TRADE treatment (far right) was carried out in a well plate assay. Representative macroscopic images of 2 week old seedlings (**j**). scale bar: 1 mm. Representative leaf vasculature images of cotyledons from plants in (**j**). Upper panel, bright field microscopy; lower panel, epifluorescence microscopy; scale bar, 50 μm. Yellow asterisks, bundle sheath cells; red asterisks, bundle sheath cells transdifferentiated into xylem cells.

**Movie 1. TRADE-GFP shows nucleo-cytosolic localisation when expressed in Arabidopsis.** Animated CLSM z-stack of *TRADE-GFP* expressing plants. Images were captured from the abaxial to adaxial side of leaves. Z-stack was 81 µm thick with 1 µm steps.

**Movie 2. TRADE-GFP localises to granular structures in bundle sheath cells** Animated CLSM z-stack of bundle sheath cells in *TRADE-GFP* expressing plants. Images were captured from the abaxial to adaxial side of leaves. Z-stack was 43 µm thick with 1 µm steps.

**Extended Data Fig. 5.**
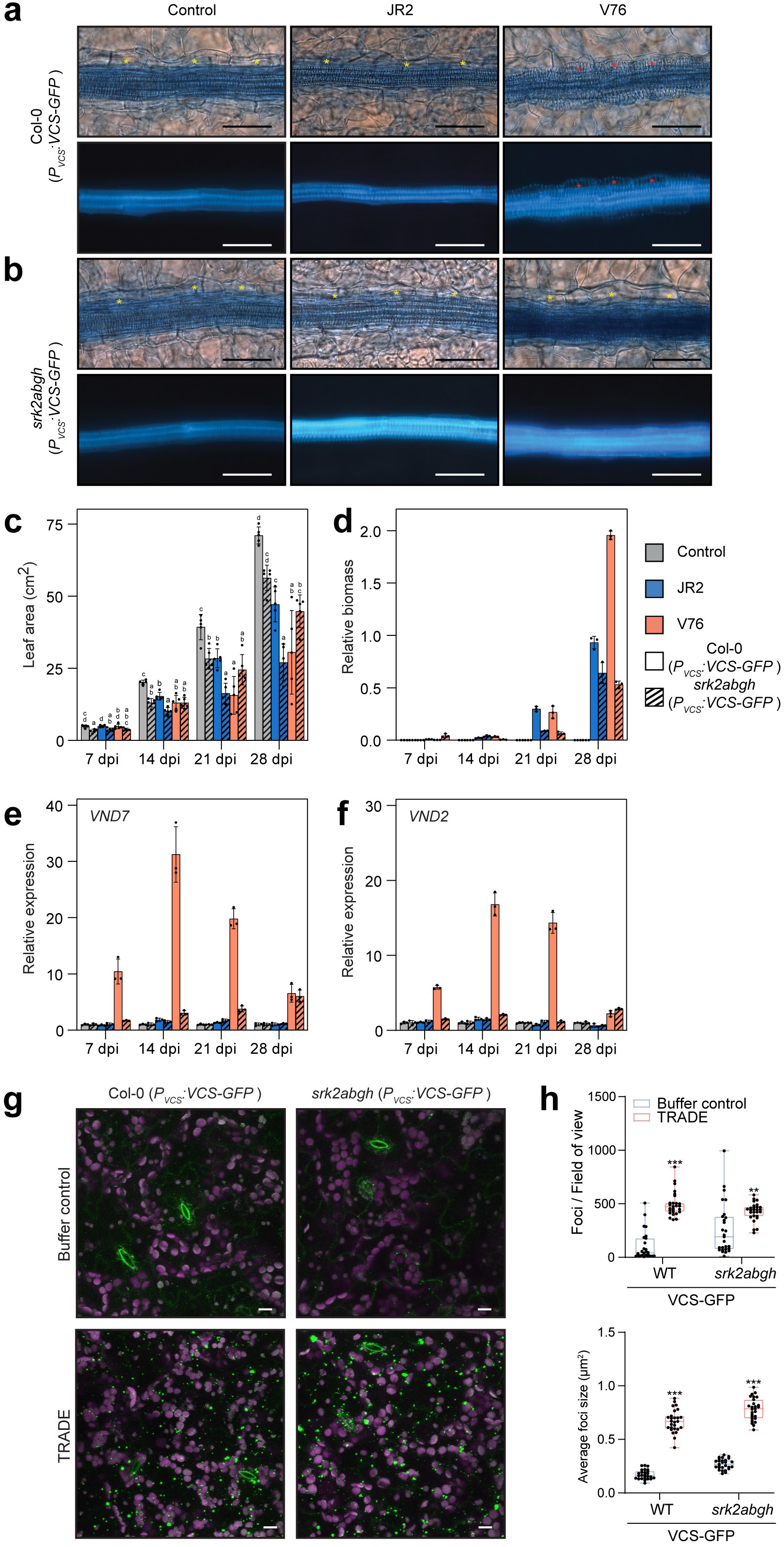
Lack of TRADE-dependent transdifferentiation and associated phenotypes in *srk2abgh* mutant plants with the exception of VCS P-body localization. **a**, **b**, Images of vasculature from control and infected *VCS-GFP* expressing plants in the (**a**) Col-0 and (**b**) *srk2abgh* backgrounds. Top row, bright field microscopy; bottom row, epifluorescence microscopy. Yellow asterisks, bundle sheath cells; red asterisks, bundle sheath cells transdifferentiated into xylem. Scale bar, 50 μm. **c**, Leaf area of control and infected *VCS-GFP* expressing plants in the Col-0 and *srk2abgh* backgrounds. Bars represent means ± SD. Letters denote significant difference within the time points (One-way ANOVA with Tukey’s HSD, p<0.05, n = 5 plants). **d**, *Verticillium* proliferation in control and infected *VCS-GFP* expressing plants in the Col-0 and *srk2abgh* backgrounds as determined with qPCR. Bars represent means ± SD of 3 technical replicates from a pool of 5 plants. **e**, **f**, Expression analysis of *VND7* (**e**) and *VND2* (**f**) in leaves of control and infected *VCS-GFP* expressing plants in the Col-0 and *srk2abgh* backgrounds. Bars represent means ± SD of 3 technical replicates from a pool of 5 plants. **g**, Maximum z-projections of CLSM images from plants expressing *VCS-GFP* in the Col-0 and *srk2abgh* backgrounds after buffer control and TRADE treatment. **h**, Quantification of foci per field of view (top) and average foci size (bottom) from plants treated in (**g**). Boxplots are represented as described in Fig. 5g. (n = 26-27, student’s t-test, **, p<0.01; ***, p<0.001). Chloroplast background fluorescence indicated as magenta in images. Scale bar, 10 μm.

**Extended Data Fig. 6.**
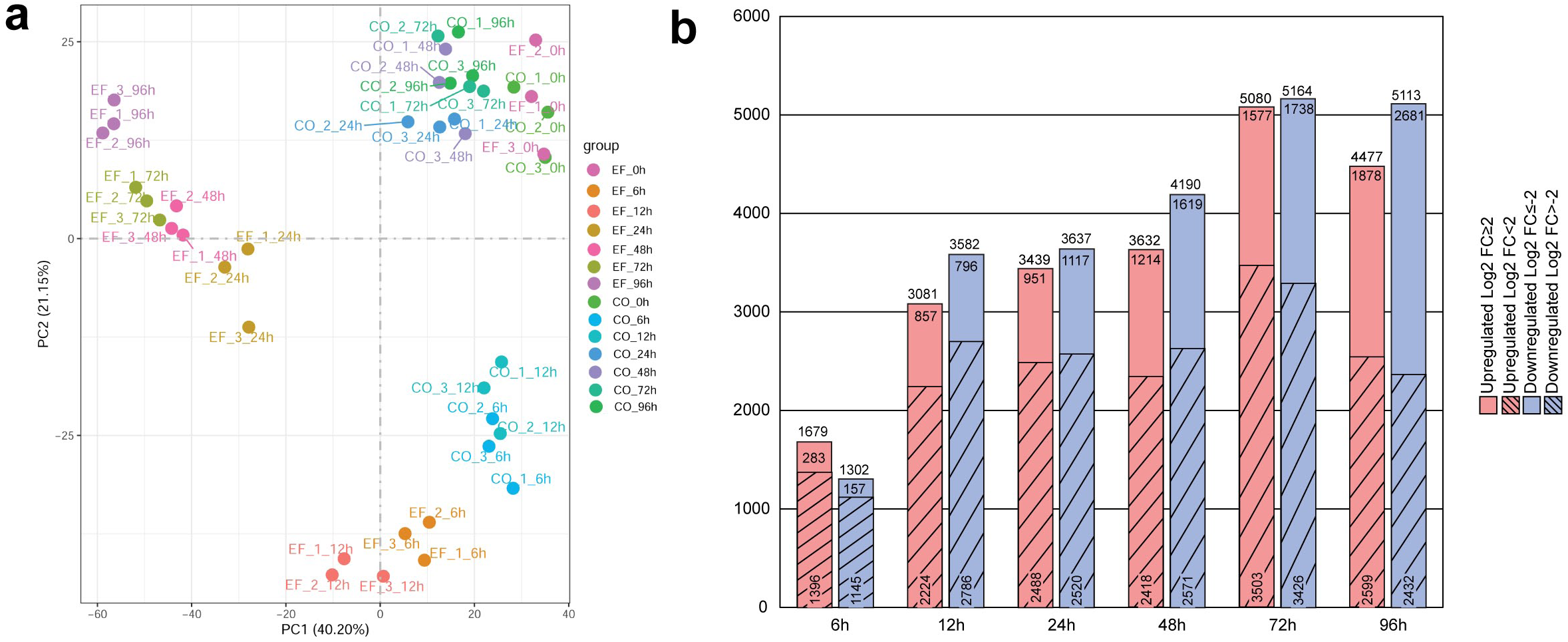
Principal component analysis (PCA) of TRADE-induced transcriptome changes and total DEGs in response to TRADE treatment. **a**, Principle component analysis (PCA) of normalized RNAseq data generated from buffer control (CO) and TRADE (EF) infiltrated samples after 0h, 6h, 12h, 24h, 48h, 72h and 96h. **b**, Total numbers of significantly upregulated and downregulated genes for time points 6-96h after TRADE treatment (padj<0.05). Counts within and on top of bars represent number of genes in each category and in total for each time point, respectively.

**Extended Data Fig. 7.**
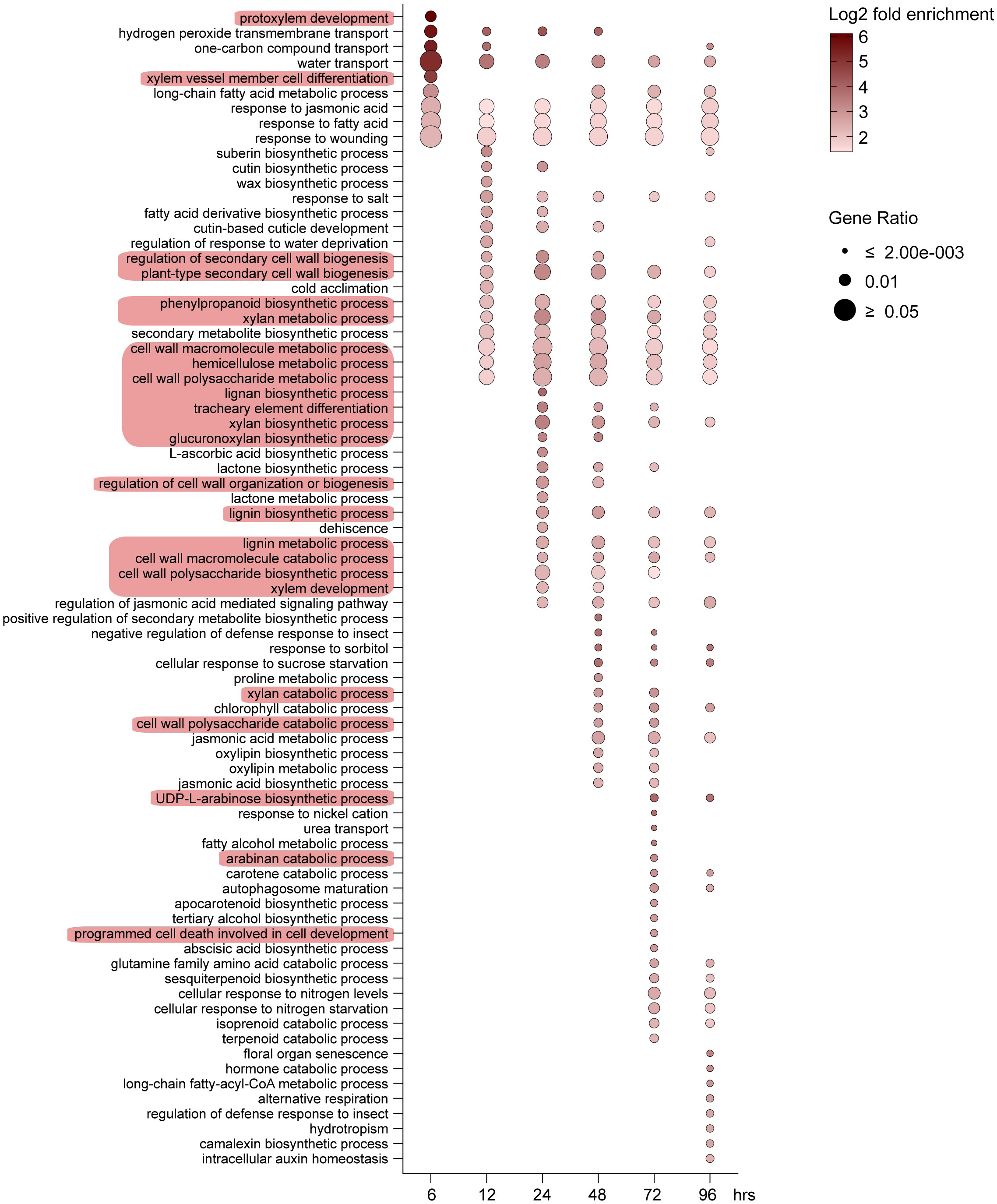
GO term enrichment analysis for upregulated DEGs at time points 6-96h. DEGs used had a log2 fold change of ≥2 (padj<0.05). Red background indicates GO terms associated with lignin, carbohydrate and cell wall biosynthesis as well as xylem/tracheary element development and differentiation.

**Extended Data Fig. 8.**
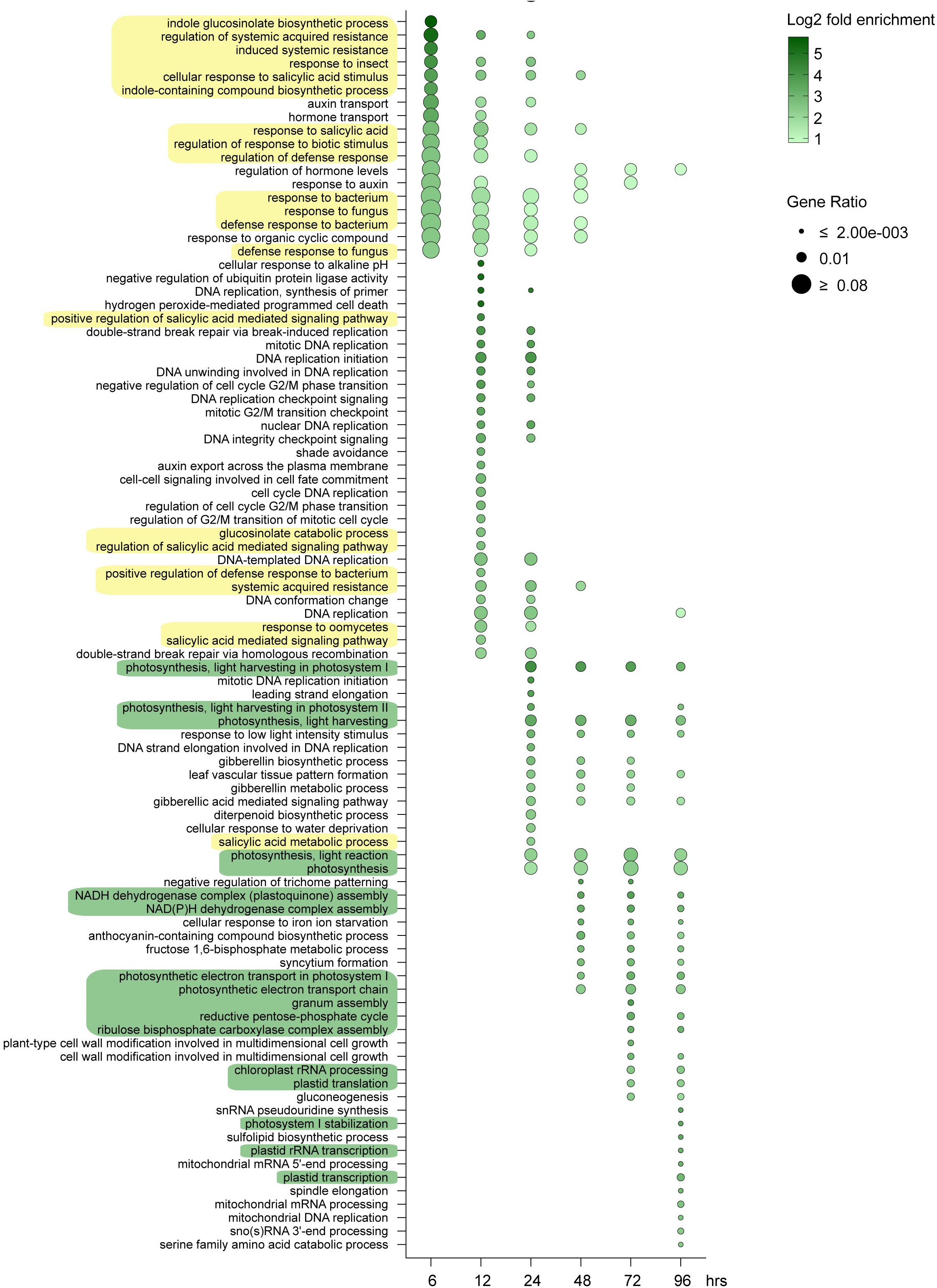
GO term enrichment analysis for downregulated DEGs at time points 6-96h. DEGs used had a log2 fold change of ≤-2 (padj<0.05). Yellow or green background indicates GO terms associated with pathogen defense or photosynthesis respectively.

**Extended Data Fig. 9.**
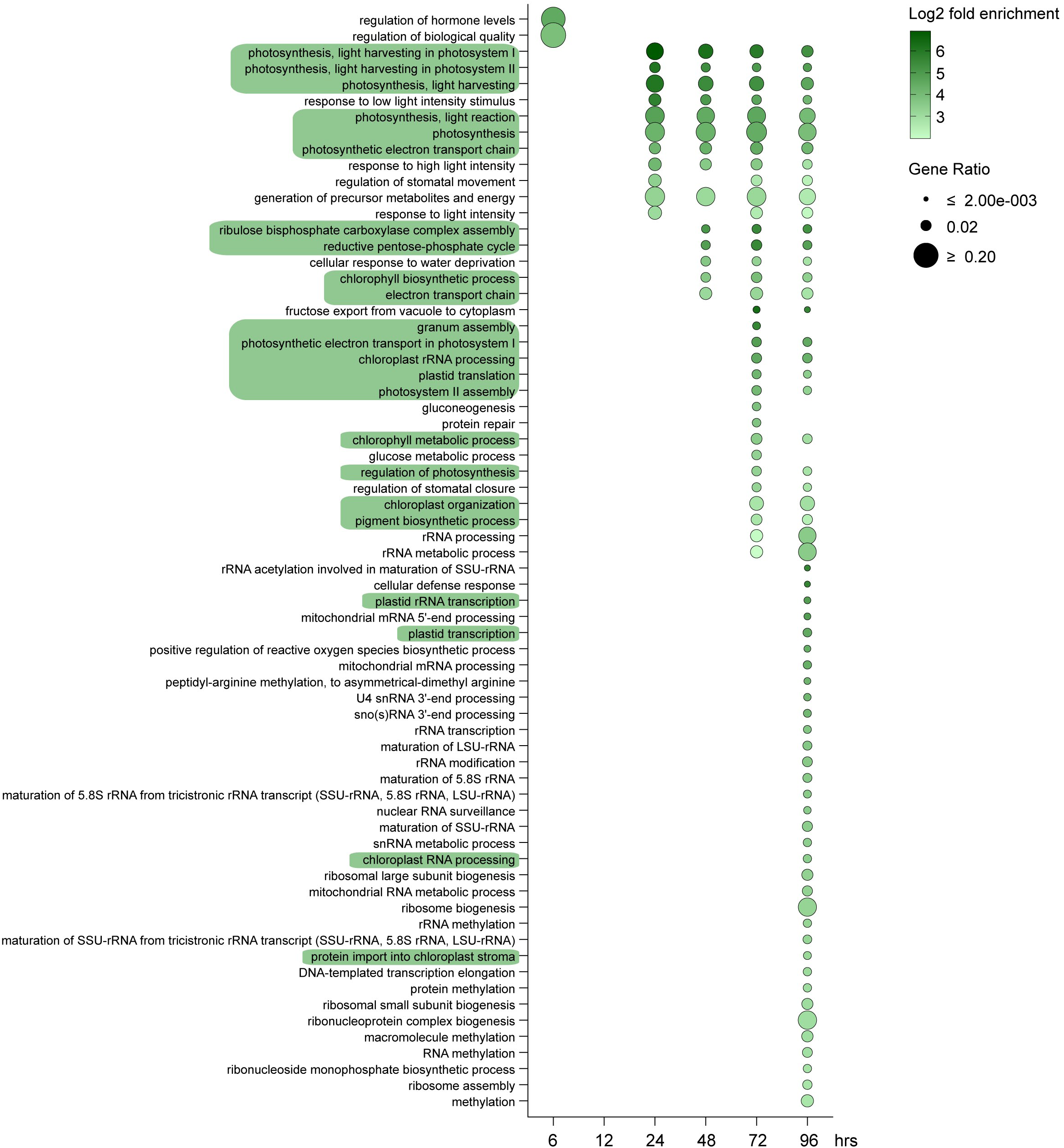
GO term enrichment analysis of downregulated DEGs overlapping between the TRADE transcriptome and the *vcs7* mutant. DEGs used had a log2 fold change of ≤-2 (padj<0.05). Green background indicates GO terms associated with photosynthesis.

## Notes

### Competing Interest Statement

The authors have declared no competing interest.

